# Gene expression is encoded in all parts of a co-evolving interacting gene regulatory structure

**DOI:** 10.1101/792531

**Authors:** Jan Zrimec, Filip Buric, Azam Sheikh Muhammad, Rhongzen Chen, Vilhelm Verendel, Mats Töpel, Aleksej Zelezniak

## Abstract

Understanding the genetic regulatory code that governs gene expression is a primary, yet challenging aspiration in molecular biology that opens up possibilities to cure human diseases and solve biotechnology problems. However, the fundamental question of how each of the individual coding and non-coding regions of the gene regulatory structure interact and contribute to the mRNA expression levels remains unanswered. Considering that all the information for gene expression regulation is already present in living cells, here we applied deep learning on over 20,000 mRNA datasets in 7 model organisms ranging from bacteria to Human. We show that in all organisms, mRNA abundance can be predicted directly from the DNA sequence with high accuracy, demonstrating that up to 82% of the variation of gene expression levels is encoded in the gene regulatory structure. Coding and non-coding regions carry both overlapping and orthogonal information and additively contribute to gene expression levels. By searching for DNA regulatory motifs present across the whole gene regulatory structure, we discover that motif interactions can regulate gene expression levels in a range of over three orders of magnitude. The uncovered co-evolution of coding and non-coding regions challenges the current paradigm that single motifs or regions are solely responsible for gene expression levels. Instead, we show that the correct combination of all regulatory regions must be established in order to accurately control gene expression levels. Therefore, the holistic system that spans the entire gene regulatory structure is required to analyse, understand, and design any future gene expression systems.

## 1. Introduction

Gene expression governs the development, adaptation, growth, and reproduction of all living matter. Understanding its regulatory code would provide us with the means to cure diseases, including heterogeneous tumours ^1^, and to control protein production for biotechnology purposes ^2,3^. Although transcriptional regulation has been a central area of research in the past decades, with advances that enable accurate measurement of mRNA levels ranging from just a few copies to several thousand per cell ^4–7^, we still cannot quantify to what extent the DNA code determines mRNA abundance, nor understand how this information is encoded in the DNA. Lack of such quantitative understanding hinders the potential of accurately controlling mRNA and protein levels by simply manipulating the sequence of the four DNA nucleotides.

A strong agreement between protein and mRNA levels in multiple organisms shows that up to 90% of the variation of protein abundance is attributed to mRNA transcription rather than protein translation ^4,5,7,8^. mRNA transcription is controlled via the gene regulatory structure, comprised of coding and *cis*-regulatory regions that include promoters, untranslated regions (UTRs), and terminators each encoding a significant amount of information related to mRNA levels ^9^. For instance, in *Saccharomyces cerevisiae*, properties of individual *cis*-regulatory regions can explain up to half of the variation in mRNA levels ^10–15^. Considering that each part of the gene structure controls a specific process related to mRNA synthesis and decay ^9,16,17^, as well as the overall transcription efficiency ^18^, all gene parts must be concerted in perfectly timed execution in order to regulate expression. However, despite the apparent importance of non-coding regions, it remains unclear how they cooperatively regulate gene expression.

Much of the current knowledge on quantitative regulation of gene expression is based on high-throughput screens of thousands of synthetic sequences studied in isolation from their native gene regulatory structures ^19–21^. These screens introduce random mutations in a non-coding region of DNA, in order to determine the effects of the mutations on gene expression. Although a *de facto* standard for expression tuning in synthetic biology, these techniques are (i) laborious and require expensive and highly-sophisticated equipment ^22^, (ii) are biased towards specificities of mutagenesis ^23^, and (iii) are often restricted to particular experimental conditions. The major problem, however, is that the biological sequence space is so large that it cannot be explored experimentally or computationally ^24^. For instance, to analyze all the possible combinations of the four nucleotides in a 20 bp promoter would require iterating over a trillion (4^20^) synthetic sequences. This limits the experimental studies to individual regulatory gene parts in the context of single reporter genes. Similarly, with natural systems, the majority of studies on mRNA transcription in the context of transcription factor (TF) binding ^25^, chromatin accessibility ^26,27^ and Chip-seq or DNase-Seq data ^28,29^, focus solely on promoter regions ^30^. Therefore, both the current natural and synthetic approaches are fundamentally limited in their ability to study the relationship between the different parts of the gene regulatory structure and their cooperative regulation of expression.

Here, we consider that DNA sequences of all living systems, through evolution, have been fine-tuned to control gene expression levels. To learn from the natural systems, we analysed 100,000 native gene sequences in over 20,000 RNA-Seq experiments from seven model organisms, including *Homo sapiens, Saccharomyces cerevisiae*, and *Escherichia coli*. In the yeast *S. cerevisiae*, the variation of gene expression per gene across the entire repertoire of different experimental conditions was 340 times lower than the variation of expression levels across all genes. Based on this high signal-to-noise ratio, our deep neural networks learned to predict gene expression levels directly from the native DNA sequences, without the need for screening experiments using synthetic DNA. Prediction of gene expression levels was highly accurate in all model organisms, and in *S. cerevisiae* (*R*^*2*^_*test*_ = 0.82) showed strong agreement with fluorescence measurements from independent published experiments. We demonstrated that, in both eukaryotes and prokaryotes, mRNA levels are determined not merely by individual coding and *cis*-regulatory regions but rather additively, by the entire gene regulatory structure. As the coding and *cis*-regulatory regions contained both orthogonal as well as overlapping information on expression levels, the entire gene codon distribution could be predicted merely from the adjacent *cis*-regulatory regions (*R*^*2*^_*test*_ = 0.58). Further mutational analysis of orthologous genes in 14 yeast species confirmed that each gene is a co-evolving unit. Next, we reconstructed the regulatory DNA “grammar” of *S. cerevisiae* by measuring the co-occurrence of sequence motifs extracted from the deep models and present across the *cis*-regulatory regions. These motif co-occurrences were found to be highly predictive of expression levels and could differentiate the expression levels of single motifs in a range of over 3 orders of magnitude. Finally, by quantifying the variation present in all 36 million promoter-terminator combinations we observed that gene expression levels change, on average, over 10-fold in either direction of the native levels. Thus with each gene, merely exchanging one side of its regulatory regions with other natural variants unlocks an enormous potential for future gene expression engineering.

## 2. Results

### The dynamic range of gene expression levels is encoded in the DNA sequence

To explore the relationship between DNA sequence and gene expression levels, we compiled a dataset of 3025 high-quality *Saccharomyces cerevisiae* RNA-Seq experiments ^31^ covering virtually all available experimental conditions from 2365 unique studies. By sorting 4975 protein-coding genes (Methods, Table S1-1) according to their median expression levels across the experiments (expressed as TPM values, i.e. Transcripts Per Million), we observed a striking trend in the expression levels (Fig 1A): 85% of the yeast protein-coding genes varied less than 1 relative standard deviation across the RNA-Seq experiments (*RSD* = *σ*/*μ*) (Fig 1B, Table S1-2). With 79% of protein-coding genes, the expression levels per gene changed merely within a one-fold range of values in ⅔ of the RNA-Seq experiments (Fig 1A). Conversely, the dynamic range of median TPM values across all the genes spanned over 4 orders of magnitude. The variance of expression levels within the whole genome was on average 340 times higher than the variance per gene across the experiments (Fig 1C). The most variable genes across the entire range of biological conditions (Fig 1B: *RSD* > 1) were significantly (Hypergeometric test BH adj. *p*-value < 0.05) enriched in metabolic processes, sporulation, and cell cycle (Fig S1-2), while the most stable genes (*RSD* < 1) were significantly (Fisher’s exact test *p*-value < 1e-16) enriched in TFIID-type constitutive promoters ^26^.

**Figure 1.**
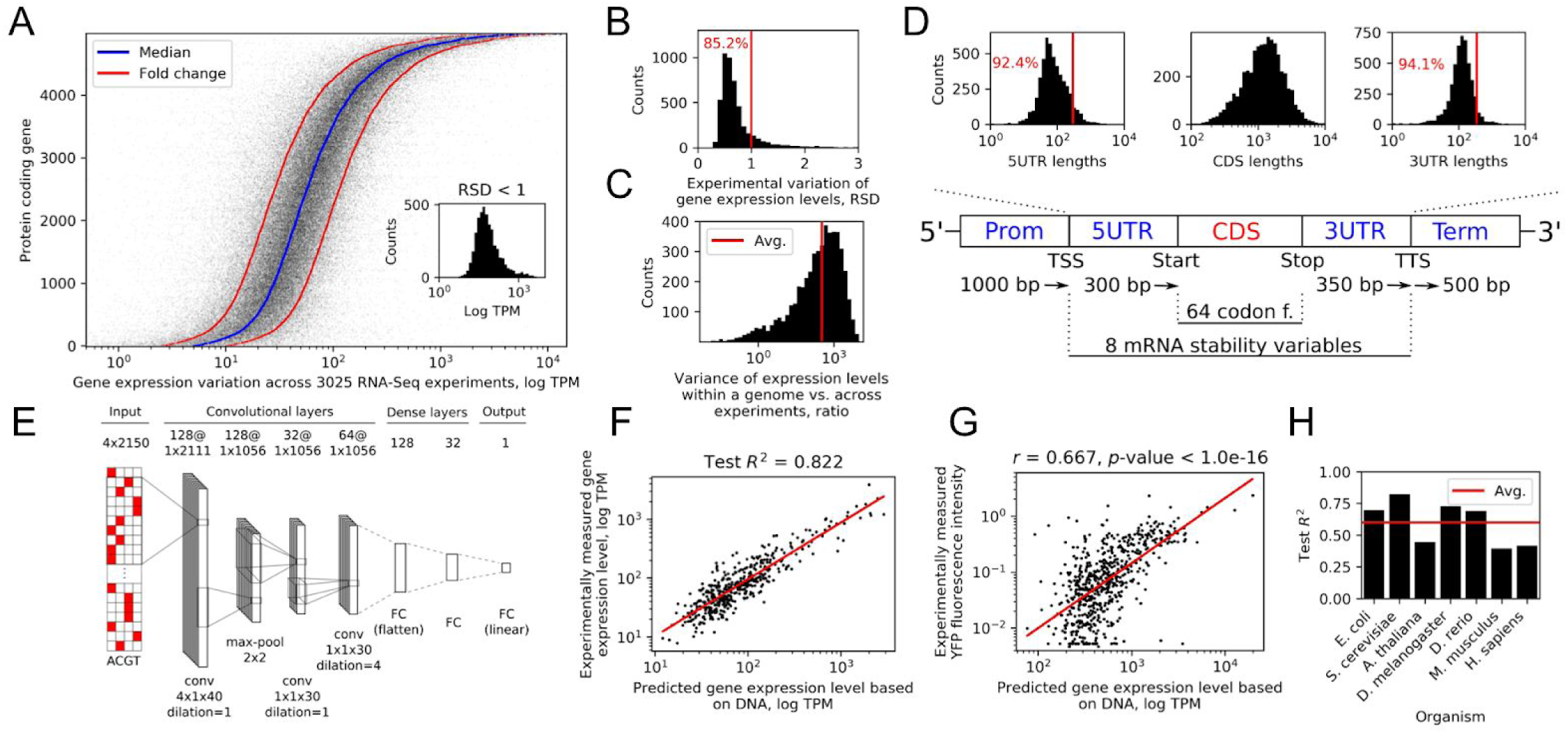
The dynamic range of gene expression levels is encoded in the DNA sequence. All results shown are with *S. cerevisiae*, except with figure (H). (A) Expression levels of protein-coding genes across 3025 RNA-seq experiments. Inset: distribution of genes with a relative standard deviation (*RSD* = *σ/μ*) below 1. (B) Experimental variation of the gene expression levels expressed as *RSD*. (C) Distribution of ratios between the variance of expression levels within a genome and the variance of expression levels per gene across the experiments. (D) Schematic diagram of the explanatory variables used for modeling, with distributions of the sequence lengths in the regions where the lengths varied. “TSS” denotes the transcription start site and “TTS” the transcription termination site. (E) The optimized deep neural network (NN) architecture, where “conv” denotes convolutional NNs, “FC” fully connected layers, and “max-pool” max-pooling layers. The values denoting sizes of parameters *layers, stride, kernels, filters, max-pooling*, and *dilation* are specified. (F) Experimentally determined versus predicted expression levels with the *S. cerevisiae* model. Red line denotes least squares fit. (G) Experimental fluorescence measurements ^39^ versus predicted expression levels. Red line denotes least squares fit. (H) *R*^*2*^ on test dataset across 7 model organisms.

To test if the observed dynamic range of gene expression levels is encoded in the DNA, we extracted the DNA sequence of the regulatory and coding regions of all the genes within one relative standard deviation of expression variation (Fig 1B: 4238 genes with *RSD* < 1). A total of 2150 bp of regulatory sequences ^9,11,14,15,32–36^, 64 codon frequencies from coding regions ^37^ and additional 8 mRNA stability variables ^13^, all known to be important for expression regulation, were used for prediction of mRNA levels (Fig 1D, Fig S1-1, Methods). Using the DNA sequence information as input, we built a regression model based on deep convolutional neural networks (CNNs) and trained it to predict the median gene expression levels (Fig 1E, Methods). The median value was a good estimator of the expression differences between the genes, as it was strongly correlated with the first principal component of the entire expression matrix (Pearson’s *r* = 0.990, *p*-value < 2e-16, Fig S1-3). To avoid potential technical biases related to read-based sequencing ^38^, mRNA levels were corrected for gene length bias (Fig S1-4, Methods). Overall, a total of 3433 gene sequences were used for training the model, 381 for tuning the model hyperparameters and 424 for testing. After optimizing the model (Methods), its predictive performance on the test set (*R*^*2*^*_test_* = 0.822, Fig 1F, Table S1-3) demonstrated that the DNA encodes the majority of the information about mRNA expression levels.

To validate the trained model on independent data, we used two published experimental datasets, where the effects of either promoter ^39^ or terminator ^40^ sequences were measured in synthetic constructs combined with fluorescence reporters. For both studies, we only inferred expression levels based on the deep neural network trained on natural genomic sequences (Fig 1E,F, Methods), meaning that the model was not exposed to the data from these studies in its training phase. The first dataset comprised measured activities of ∼900 native yeast promoters recorded in synthetic constructs with a single strong terminator (ADH1) and a YFP fluorescence reporter in 10 different conditions ^39^. In all 10 conditions, the predictions of mRNA levels inferred based on the DNA sequences of the synthetic constructs and YFP codon frequencies were in strong agreement (Pearson’s *r* from 0.570 up to 0.718, *p*-value < 1e-16) with the experimental YFP readouts (Fig 1G: median YFP readout shown, Fig S1-5). Similarly, the second experimental dataset ^40^ contained expression measurements of over 5000 terminators with a fixed strong promoter (TDH3) and a GFP fluorescence reporter. Despite the fact that predictions based on these synthetic constructs were only moderately correlated (Pearson’s *r* = 0.310, p < 1e-16) with the measured protein fluorescence intensities (Fig S1-5), this result was in fact stronger than the correlation between reporter fluorescence and measured mRNA abundances reported in the original study (Pearson’s *r* = 0.241) ^40^.

Equally as with yeast, we processed an additional 18,098 RNA-Seq experiments from 1 prokaryotic and 5 eukaryotic model organisms (Table S1-1, Methods), which displayed similar properties as yeast (Table S1-2). The organisms were selected to cover the whole known range of genome regulatory complexity, from 892 genes/Mbp (*Escherichia coli*) to 6 genes/Mbp (*Homo sapiens*), which is known to affect the gene structure and regulation ^41^. After training the models for each organism, the predictive power on test data (*R*^*2*^_*test*_) varied from 0.394 for *Mus musculus* to 0.725 for *Drosophila melanogaster* (Fig 1H, Table S1-3). Overall, the predictions were less accurate for higher eukaryotes, which could be attributed to the increase in transcriptional complexity, e.g. due to alternative splicing ^42^, expression differences across tissues ^43^, and distant enhancer interactions ^44–47^, which were not accounted for in the present models. The prediction performance was thus correlated (Pearson’s *r* = 0.616, *p*-value < 3e-3) with the genomic complexity of the model organisms (Fig S1-6). Nevertheless, the average performance across model organisms (*R*^*2*^*_test_*) of 0.6 (Fig 1H) indicated that the majority of mRNA expression differences in all organisms can be predicted directly from the DNA.

### Coding and cis-regulatory regions additively contribute to gene expression prediction

To evaluate the importance of each part of the gene regulatory structure (Fig 1D) for the prediction of expression levels, we measured the amount of relevant information in each regulatory region. Similarly to the complete model (Fig 1E,F), we trained multiple CNN models independently on promoter, 5’-UTR, 3’-UTR, and terminator regions as well as their combinations. To justify the use of deep convolutional networks, as a baseline we performed shallow modeling using a variety of regression algorithms, including multiple linear regression, elastic net, random forest, and support vector machines with nested cross-validation (Fig 2A, Methods). The single regulatory regions alone could explain less than 28% of variation in mRNA abundance levels, which suggested that the entire gene structure was important for controlling gene expression levels (Fig 2B, Fig S2-1). Each DNA region thus additively contributed to the prediction of mRNA levels and increased model performance, with the model trained on all four regulatory regions accounting for approximately 50% of the mRNA abundance variation (*R*^*2*^*_test_*= 0.492, Fig 2A,B, Table S2-1). In contrast, none of the shallow models could predict gene expression levels from the regulatory sequences (*R*^*2*^*_test_* < 0.031, Fig 2A, Table S2-2). Despite the observation that the sole mRNA stability variables were also somewhat informative about gene expression levels (*R*^*2*^*_test_* = 0.378, Fig 2A), they did not improve the overall model performance next to the codon frequencies and regulatory sequences, likely due to their information redundancy with these variables (Figure S2-2: *R*^*2*^*_test_* was 0.779 when predicting mRNA stability variables using regulatory sequences).

**Figure 2.**
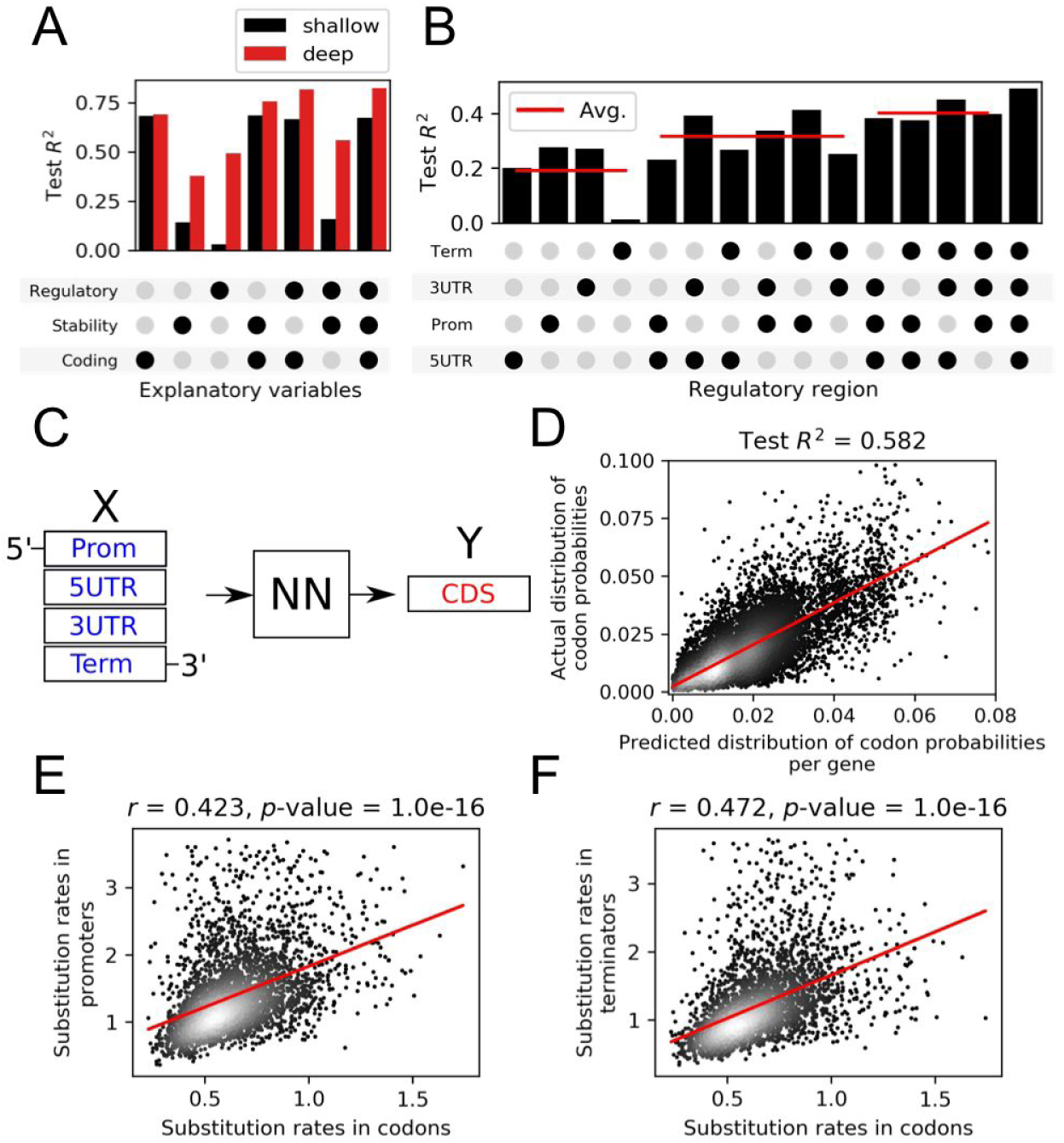
Coding and cis-regulatory regions additively contribute to gene expression prediction. (A) Test *R*^*2*^ with different combinations of *cis*-regulatory regions and deep modeling. (B) Test *R*^*2*^ with different combinations of coding (codon prob.), mRNA stability (see Fig S1-1C) and *cis*-regulatory regions, using shallow and deep modeling. (C) Schematic depiction of the regulatory regions as input, and coding regions (codon prob.) as target variables with deep neural networks (NN). (D) Actual versus predicted distribution of codon probabilities obtained with models obtained using strategy in (C). Red line denotes least squares fit. (E) Evolutionary substitution rates in promoter vs. coding regions in orthologous genes of 14 yeast species. Red line denotes least squares fit. (F) Evolutionary substitution rates in terminator vs. coding regions in orthologous genes of 14 yeast species. Red line denotes least squares fit.

On the other hand, the codon frequencies alone could explain well over 50% of mRNA level variation both with shallow and deep models (*R*^*2*^_*test*_ was 0.681 and 0.690, respectively, Fig 2A, Table S2-2). Considering that the amount of information was comparable to, or greater than that of the regulatory sequences, we attempted to measure the amount of information overlap in these regions. We thus trained a CNN to predict the entire codon frequency vector for each gene using only its regulatory regions (Fig 2C, Methods), which showed that over 58% of the gene’s codon frequency variation was directly encoded in its adjacent regulatory regions (Fig 2D: *R*^*2*^_*test*_ = 0.582). This result suggested that the coding and noncoding regulatory regions might have co-evolved under common evolutionary pressure. To test this hypothesis, we composed a dataset of cis-regulatory regions of orthologous genes from a diverse set of 14 yeast species ^48^ (Table S2-3) and compared the mutation rates between the different regions of the yeast gene structure (Methods). The mutation rates of the promoter and terminator regions displayed a moderate positive correlation (Pearson’s *r* = 0.423 and 0.471, *p*-value < 1e-16, respectively) with the mutation rates of the yeast coding regions (Fig 2E,F), supporting the co-evolution hypothesis.

### Deep learning identifies specific DNA positions controlling gene expression levels

To explore the information learned by the deep neural network (Fig 1E,F) and identify the specific parts of the DNA sequences that were most predictive of gene expression levels, we developed a pipeline for evaluating the *relevance* of each specific DNA sequence position in relation to the predicted gene expression levels (Fig S3-1, Methods). Briefly, we removed sliding windows of 10 base pairs (Fig S3-2) along each gene’s regulatory sequence and compared the model’s response to the occluded sequences with predictions on the original unmodified DNA ^49,50^. The occluded parts of the input DNA sequences that significantly deviated (exceeding ± 2 standard deviations) from the original data were regarded as the most relevant for gene expression changes (Fig 3A). The largest density of relevant regions was obtained in direct vicinity of the boundary sites defining regulatory and coding sequences, to which the sequence data were anchored (see Fig 1D). On average 214 base pairs (bp) in promoters, 74 bp in 5’-UTRs, 94 bp in 3’-UTRs, and 127 bp in terminator regions per gene significantly affected the prediction of gene expression levels (Fig 3A). *Relevance* profiles of promoter regions were also strongly correlated (Pearson’s *r* = - 0.7, *p*-value < 1e-16) with experimentally measured nucleosome occupancy scores ^27^ (Fig S3-3), which suggested that the deep learning algorithms uncovered the intrinsic molecular information encoded in the nucleotide composition.

**Figure 3.**
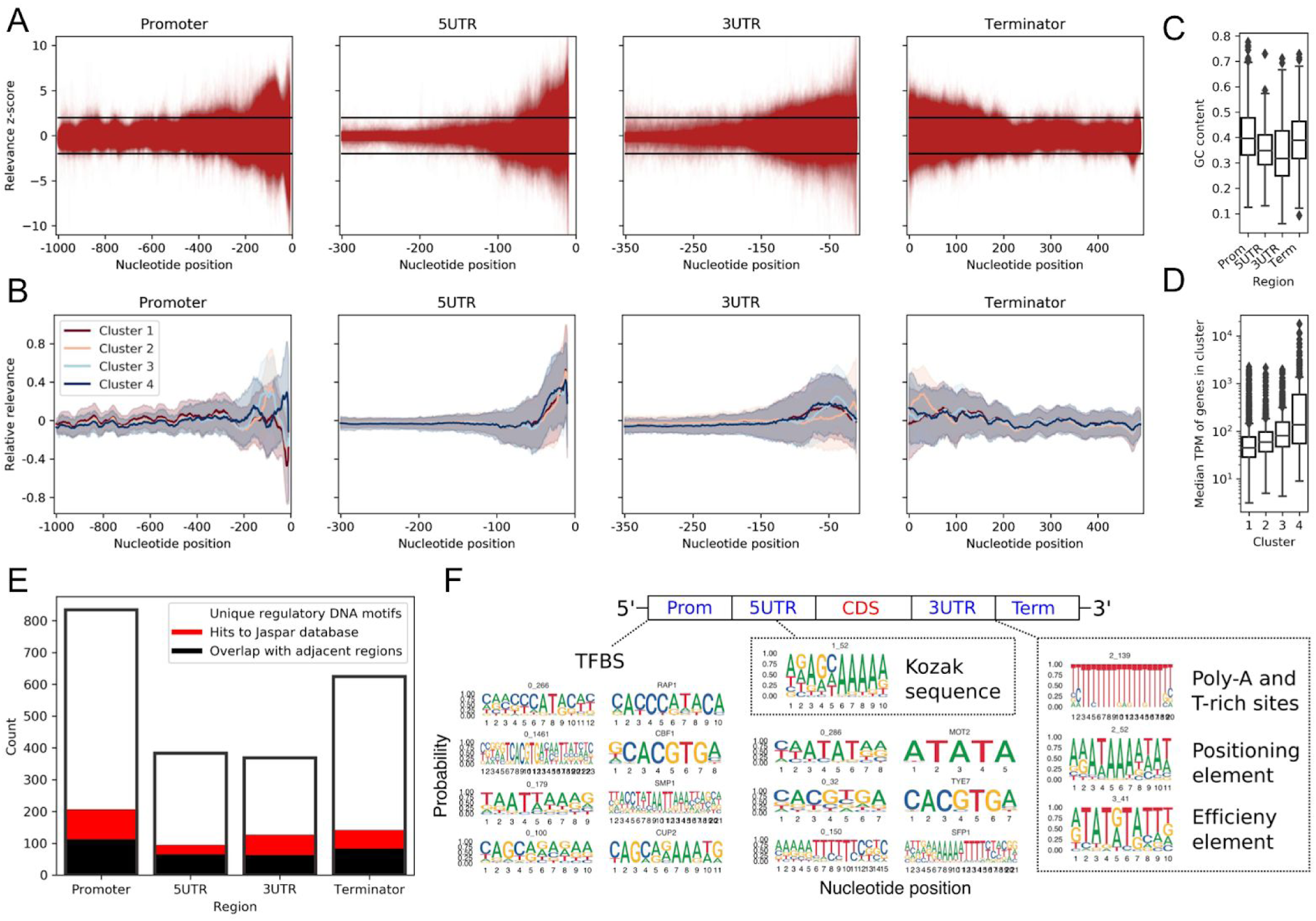
Deep learning identifies the DNA positions predictive of expression levels. (A) *Relevance* profiles across *cis*-regulatory region sequences obtained by querying the deep models (Fig 1F). The decrease in *relevance* of UTR sequences at left edges was due to the large number of sequences that were shorter than the analysed regions (see Fig 1D). (B) Clustered *relevance* profiles across the *cis*-regulatory region sequences. (C) GC content in the *cis*-regulatory regions. (D) Median expression levels of genes in the clustered *relevance* profiles. (E) Amount of regulatory DNA motifs uncovered in the *cis*-regulatory regions and amount of motif overlap between adjacent regions as well as with the Jaspar database ^53^. (F) Examples of regulatory DNA motifs uncovered across all the *cis*-regulatory regions that correspond to published motifs and sequence elements (see Table S1-1C). “TFBS” denotes transcription factor binding sites.

Clustering the gene regulatory regions according to their expression *relevance* profiles (Methods) identified 4 stable clusters that significantly (Rank sum test *p*-value < 1e-4) differed in the positional information (Fig 3B) and were significantly (Rank sum test *p*-value < 1e-16) informative about gene expression levels (Fig 3D). Cluster 4, which contained highly expressed genes, was significantly (Hypergeometric test *p*-value < 1e-10) enriched in the occupied proximal-nucleosome (OPN) regulation strategy as opposed to the depleted proximal-nucleosome (DPN) strategy ^51^, which was likely related to the concurrent enrichment (Fisher’s exact test *p*-value < 1e-12) of inducible SAGA promoters in the cluster. Cluster 4 also comprised genes with higher transcriptional plasticity spanning an over 4-fold higher variability of expression levels compared to the other clusters (Levene’s test *p*-value < 1e-16). As is typical for highly abundant proteins, such as metabolic enzymes, that are linked to less defined nucleosome positions and higher dependence on nucleosome remodelling ^51,52^, the cluster was significantly enriched (Hypergeometric test BH adj. *p*-value < 0.01) mostly in metabolic processes (Fig S3-4). In contrast, Cluster 1, with lowly expressed genes, was related (Hypergeometric test BH adj. *p*-value < 0.01) to cell cycle regulation and DNA repair (Fig S3-4). The largest differences in positional expression *relevance* were identified in promoter and terminator sequences (Fig 3B). For instance, in promoters of lowly and highly expressed genes (Clusters 1 and 4, respectively), occluding the original sequences yielded opposite effects. These positional differences were independent of the overall nucleotide composition (Fig S3-5), which indicated that specific regulatory sequences were likely responsible for defining the expression levels.

From the set of all significantly relevant DNA sequences (Fig S3-6), we identified the specific regulatory DNA motifs important for predicting expression levels using clustering and alignment (Methods). The highest quality motifs were obtained with the 80% sequence identity cutoff, according to the following criteria: (i) genome coverage, (ii) the amount of retained relevant sequences in motifs (seq. coverage) and (iii) % overlap with known motifs in databases (Table S3-1). Over 2200 expression related regulatory DNA motifs were uncovered across all 4 regulatory regions (Fig 3E). The majority of motifs were unique to each specific region, as analysis of motif similarity across the adjacent regulatory regions (Methods) showed that, on average, less than 16% of the motifs significantly (BH adj. *p*-value < 0.05) overlapped between the regions (Fig 3E, Table S3-2). This further supported our observation that every regulatory region contains additional information related to the gene expression levels (see Fig 2A). Further comparison to JASPAR ^53^ and Yeastract ^54^ databases (Methods) showed that on average, approximately 13% of the identified motifs (Fig 3E) were significantly (BH adj. *p*-value < 0.05) similar, 38% and 63% respectively, to the known transcription factor binding sites (TFBS) recorded in these databases (Fig S3-7). The majority of these motifs were identified not only in the promoter but also in the terminator region (Fig S3-7), due to the overlap between neighboring genes ^55^. A significant (Rank sum test *p*-value < 1e-8) decrease in GC content in the UTR regions (Fig 3C) indicated that the identified motifs were likely to contain the regulatory DNA signals from UTR regions and terminators (Fig 3F, Fig S1-1). This included the 5’-UTR Kozak sequences and 3’-UTR processing DNA elements that are enriched in the thermodynamically less stable A/T nucleotides ^11,14,34,56,57^. The deep models therefore successfully identified known regulatory signals (Fig 3F) as well as uncovered new regulatory signals across all the gene regulatory regions.

### Motif co-occurrence uncovers the regulatory rules of gene expression

To determine the regulatory meaning of the DNA motifs reconstructed from the deep learning *relevance* profiles (Fig 3F), we analysed their informative power relative to the specific gene expression levels. For this, we calculated the signal-to-noise ratio (*SNR* = *μ*/*σ*) of expression levels across genes that carried the identified motifs (Fig 4A, Fig S4-1). The spread of expression levels of motif-associated genes was over 2.5-fold larger than its expected (median) expression level (Fig S4-2), resulting in a low median *SNR* of under 0.4 (Fig 4A). Of the full 4 order-of-magnitude range of gene expression levels observed in the data (Fig 1A), only 57% could be recovered with the motifs, as indicated by the average expression level of genes per associated motif (Fig 4B, Fig S4-1B). The range of gene expression levels for the identified motifs were thus too dispersed and overlapped, indicating that the predictions made by the model (Fig 1F, Fig 3B) were likely not based on single motif occurrences.

**Figure 4.**
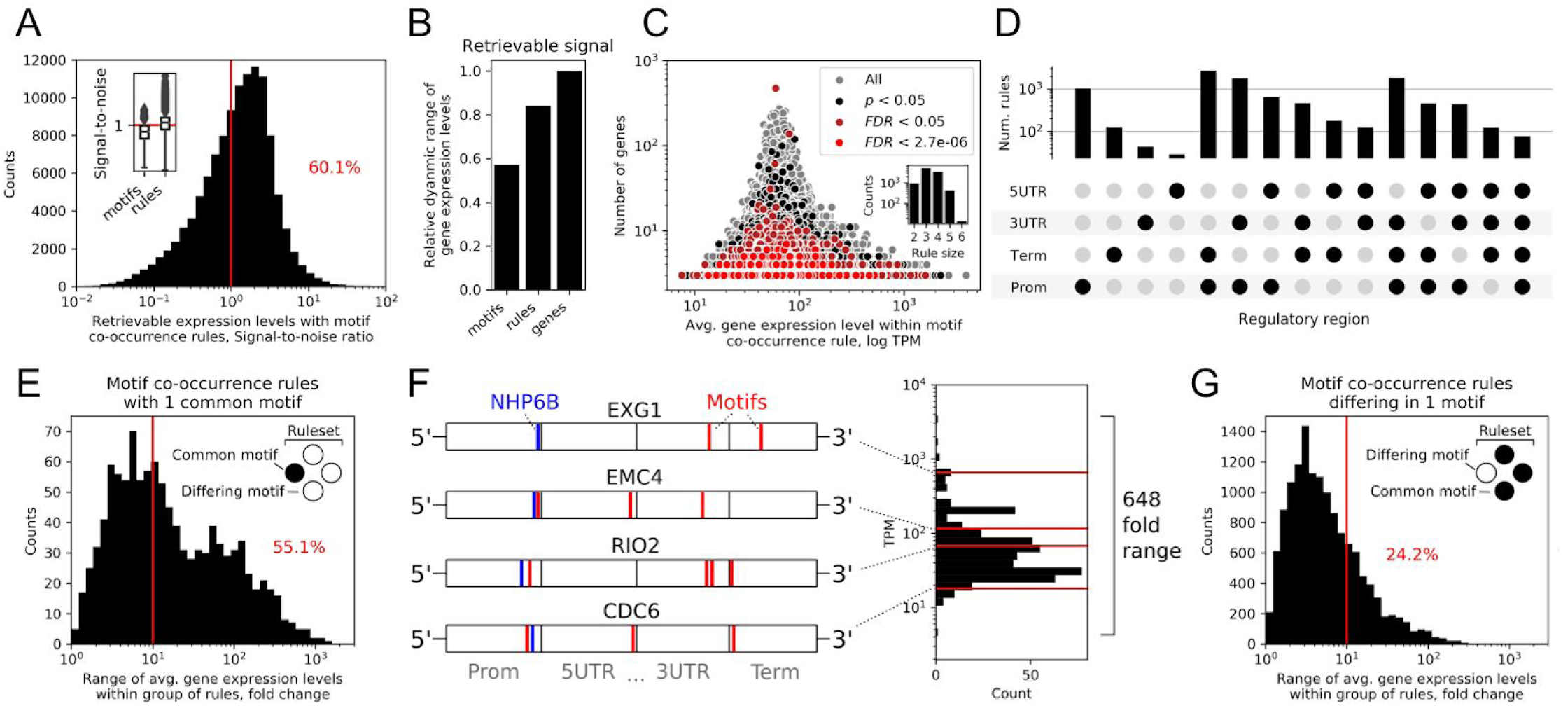
Motif co-occurrence uncovers the regulatory rules of gene expression. (A) Distribution of the signal-to-noise ratio (*SNR*) of expression levels across genes carrying the identified motif co-occurrence rules. Inset: comparison of *SNR* with single motifs and motif co-occurrence rules. (B) The amount of the full observed range of gene expression levels that could be retrieved by the average expression level across genes defined either by single motifs or motif co-occurrence rules. (C) The amount of genes in a motif co-occurrence rule versus the average expression level across genes, defined by that rule, with increasing statistical significance levels. Inset: distribution of the number of co-occurring motifs in significant (*FDR* < 0.05) rules. (D) Distribution of motif co-occurrence rules across single or multiple *cis*-regulatory regions, according to the locations of the co-occurring motifs. (E) Distribution of gene expression levels with groups of motif co-occurrence rules that have one rule in common (ie. unchanged). (F) Illustration of four genes (CDC6, RIO2, NSP1, EXG1) common to a group of motif co-occurrence rules that span a 648-fold range of gene expression levels. Common to all these rules is a NHP6B transcription factor binding site (BH adj. *p*-value < 0.005, SGD:S000002157 ^59^) in their promoter (blue line), whereas all diverge in possessing 2 to 4 other DNA motifs (red lines) across the whole gene regulatory structure, which define their expression levels. Red lines in the histogram denote the specific expression levels of the genes. (G) Distribution of gene expression levels with groups of motif co-occurrence rules that differ by a single motif.

Alternatively, the observed interactions across the regions (Fig 2A,B) and positional differences in expression levels (Fig 3B) suggested that statistically measured interactions of motifs might carry a greater indication of expression levels than single motifs (Fig 3F). We therefore searched for patterns of co-occurring motifs, i.e. motifs that are more likely to be present in genes together with other motifs than alone, and termed them regulatory ‘rules’. Here we used market basket analysis - a technique that is commonly used for identifying frequently bought items in market research ^58^ (Methods). The number of identified rules corresponded to the quality of identified motifs, and was largest at the 80% sequence identity cutoff, decreasing markedly when increasing the identity cutoff (Table S3-1). A total of 9,962 rules were significantly overrepresented (BH adj. *p*-value < 0.05) in at least 3 genes and across 93% of the analysed protein coding genome, and spanned 62% of all unique motifs that represented 86% of all motif occurrences (Fig 4C, Fig S4-3). Importantly, the rules were frequently found in smaller groups of genes (at most 10 genes) and were crucial for discerning genes with low and high expression levels, as the dynamic range of expression values decreased by 2-fold in rules present in more than 10 genes (Fig 4C). Similarly, rules comprised of a larger number of co-occurring motifs (Fig 4C: rules of up to 6 motifs found) were also sparser and occurred in smaller groups of genes (Fig S4-4). In total, 88% of the significant rules (BH adj. *p*-value < 0.05) co-occurred across the regulatory regions, compared to the rest that were present within single regions (Fig 4E). This resulted in over 6-fold more connections between motifs in promoters, terminators, and 3’-UTR regions than within any single region (Fig 4E).

By comparing the genes that carry the single motifs to those with co-occurring motifs in rules, we observed that the range of average expression values of genes spanned by the rules exceeded those of single motifs by over 11-fold (Levene’s test *p*-value < 1e-16) (Fig S4-2) and recovered, in total, 84% of the whole range of gene expression levels (Fig 4B, Fig 1A). Furthermore, a significantly (Rank sum test *p*-value < 1e-16) narrower window of expression levels was observed with rule-associated genes compared to single motifs, as the variance of expression levels was over 16-fold lower (Fig S4-2). This shows that the genes containing co-occurring motifs fall under the precise control of the specific co-occurrence rule. The signal-to-noise ratio, which exceeded 1 for over 60% of rules, in contrast to motifs, of which 78% were below 1 (Fig 4A, Fig S4-1B), demonstrated that the precision of expression control of the rule-associated genes was, on average, 3-fold higher (Rank sum test *p*-value < 1e-16) than of single motifs (Fig S4-1). This again showed the presence of statistically measurable interactions across the entire gene regulatory structure, but this time at the level of motifs, thus supporting the existence of a coevolved interacting regulatory DNA grammar.

To analyse how differences in the motif co-occurrence context across the regulatory regions affect expression levels of genes, we grouped the motif co-occurrence rules, such that one motif was left unchanged, while the other motifs differed across the genes (Fig 4E, rules with at least 3 motifs were used). The motif co-occurrence context across the regulatory regions repurposed the common motif in a range of up to 1484-fold change of expression levels. Of the 1079 such rule sets that repurposed a given motif, 55% changed by at least one order of magnitude of expression levels (Fig 4E). The repurposed motifs included significant (BH adj. *p*-value < 0.05) hits to a range of Jaspar TFBS in promoter regions, which co-occurred with, and were repurposed by, motifs across all the *cis*-regulatory regions (Table S4-1). For example, one of the largest ranges of expression levels was observed with a group of NHP6B-like motifs (SGD:S000002157, 648-fold change), a TFBS that binds to and remodels nucleosomes. These co-occurred with motifs from the adjacent regulatory regions that govern the expression levels of many essential yeast genes (Fig 4F), including those involved in DNA replication (CDC6), ribosomal RNA processing (RIO2), and coding for nuclear (NSP1) and cell membrane (EXG1) proteins. Furthermore, a similar trend was observed by an alternative grouping of the motif co-occurrence rules, such that only one motif differed across the genes. Even a single motif substitution could achieve up to a 605-fold change of expression levels, where 24% of such rulesets changed by at least one order of magnitude of expression levels (Fig 4G). Finally, an analysis of genes sets containing the most widespread motif co-occurrence rules (that covered 20 or more genes) did not show any significant enrichment of specific cellular functions, which suggested that the uncovered gene expression grammar is shared across genes of all functions and is an integral part of the basic gene regulatory structure (Fig 1D).

Since the regulatory regions were found to be coevolved with the coding region (Fig 2E,F) and highly predictive of the codon frequencies (Fig 2D), we analysed the properties of codon frequencies across the motifs and rules similarly as with the expression levels. The range of median Euclidean distances between codon frequencies, defined by the co-occurring motifs in rules, exceeded that of the single motifs by 6.5-fold (Levene’s test *p*-value < 1e-16), whereas the average variance decreased almost 5-fold (Rank sum test *p*-value < 1e-16, Fig S4-5). This showed that, compared to the motifs, co-occurrence rules defined more conserved ranges of codon frequencies, and that the regulatory DNA grammar learned by the deep models and highly predictive of gene expression levels (Fig 1F) was based on the combined properties of regulatory as well as coding regions.

### Regulatory properties learned from native genome guide expression engineering

The common practice to manipulate gene expression levels is to combine terminator (or promoter) sequences with specific strong promoters (or terminators) ^39,40,60,61^. Considering that our findings point to a strong dependence of gene expression on the interaction between all regions of the gene regulatory structure (Fig 2A,B,D, Fig 4E,F,G), we used the deep neural networks (Fig 1E,F) to explore how much the expression level of each gene can be varied with all the possible promoter-terminator combinations, a task that would be otherwise challenging to perform experimentally. To rationally simplify this analysis and retain just two global halves of the gene regulatory structure, here 5’-UTRs were combined with promoters and 3’-UTRs with terminators (Fig 5A). For each half, 17,960,644 combinations of native gene promoters or terminators with the corresponding variable counterparts were tested using the same dataset as for training the models (Fig 1A: RSD < 1). We observed that, on average, varying the terminator region introduced a significant (Rank sum test *p*-value < 1e-16) 3-fold change in either direction of expression levels, compared to the native gene structure (Fig 5B). The terminator constructs achieved up to a 130-fold increase (Fig 5B: YOL097W-A promoter with TDH3 terminator) and 14-fold decrease (Fig 5B: TIM10 promoter with TOP3 terminator) of expression levels compared to the native counterparts, which showed that with a given gene, a range of over two orders of magnitude of expression levels (Fig 5B: 130-fold range with YOL097W-A) could be unlocked merely by exchanging the terminator region. This held true for both strong as well as weak promoters (Fig 5C), as for instance, with the commonly used strong promoter PGK1 ^60,61^, we identified a regulatory combination where the expression levels decreased 3.2-fold (Rank sum test *p*-value < 1e-16) compared to the native counterpart. Conversely, with the natively weak promoter REV1, a terminator context with a significant (Rank sum test *p*-value < 1e-16) increase of over 7-fold was identified (Fig 5C). A further comparison of natively weak and strong promoters (100 top and bottom sorted constructs) confirmed that weak ones were expressed overall 2-fold more highly (Rank sum test *p*-value < 1e-16), and strong ones overall 1.3-fold more lowly (Rank sum test *p*-value < 1e-16) than the native sequence (Fig S5-1). The computed degree of regulatory freedom was supported also by published experimental results with the TDH3 promoter ^40^ (Fig 5D). Despite the specific experimental conditions leading to an offset in the measurements, we observed a moderate correlation (Pearson’s *r* = 0.310, *p*-value < 1e-16) to the variability of the fluorescence intensities with 4005 different terminators ^40^ as well as a comparable overall dynamic range (Fig 5D).

**Figure 5.**
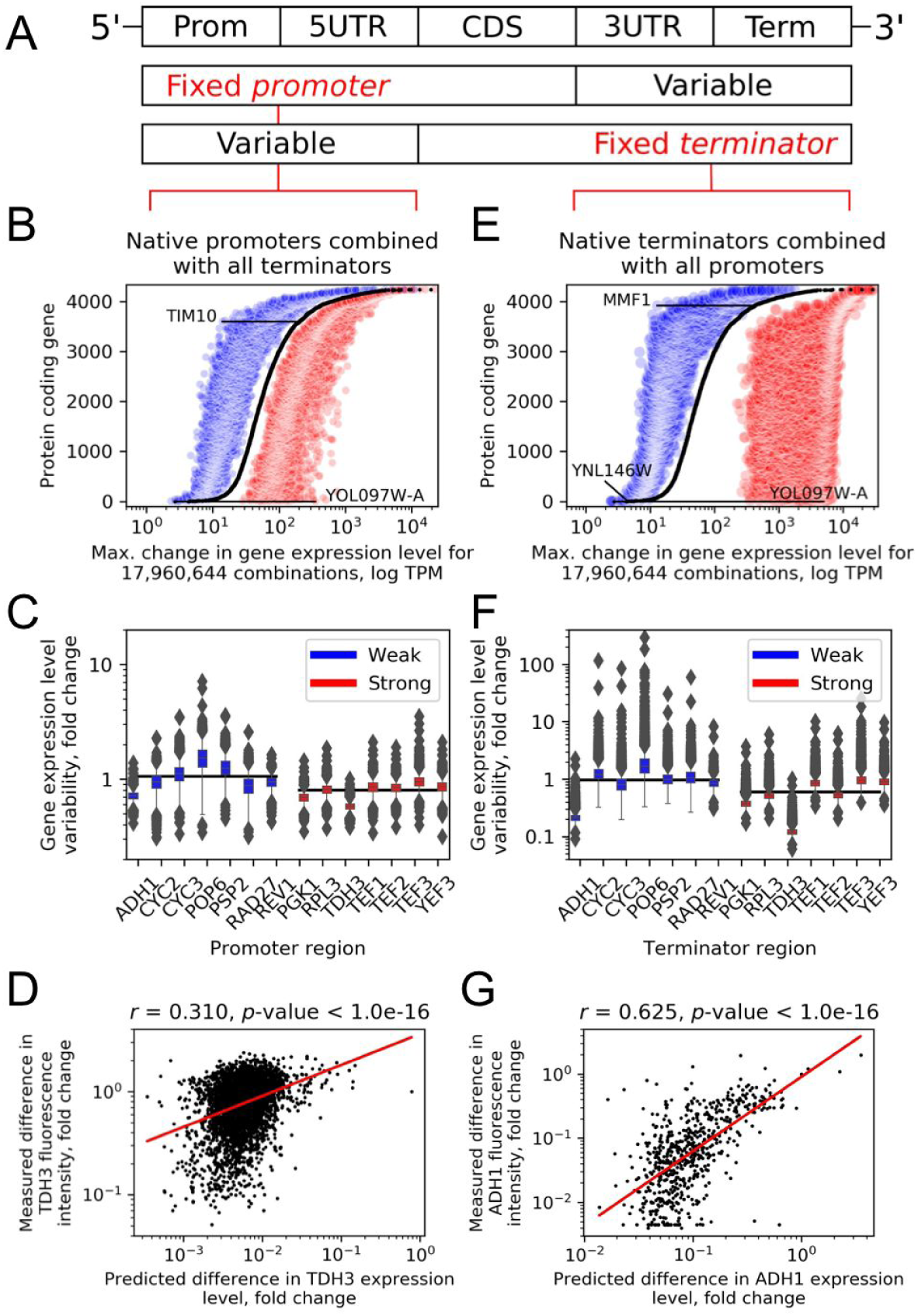
Regulatory properties learned from native genome guide expression engineering. (A) Two global halves of the gene regulatory structure where 5’-UTRs were combined with promoters and 3’-UTRs with terminators. (B) Maximum increases (red) and decreases (blue) of gene expression levels with combinations of native promoters with all terminator variants. (C) Distributions of gene expression levels for combinations of either strong or weak promoters with all terminator variants. Black lines denote median exp. levels. (D) Comparison of changes in predicted TDH3 expression levels with measured fluorescence intensities when combining the promoter with 4005 terminator variants ^40^. Red line denotes least squares fit. (E) Maximum increases (red) and decreases (blue) of gene expression levels with combinations of native terminators with all promoter variants. (F) Distributions of gene expression levels for combinations of either strong or weak terminators with all promoter variants. Black lines denote median expression levels. (G) Comparison of changes in predicted ADH1 expression levels with measured fluorescence intensities when combining the promoter with 625 terminator variants ^39^. Red line denotes least squares fit.

By performing an analogous analysis using terminator regions and instead varying the promoters, we observed, on average, an over 20-fold change (Rank sum test *p*-value < 1e-16) in either direction of expression levels and a dynamic range of over 3 orders of magnitude (Fig 5E: 2120-fold range with YNL146W). For all terminators, promoters could be identified that exerted a strong positive stabilizing effect, as the most pronounced variation was observed in the direction of increasing expression levels up to 1847-fold (Fig 5E: YOL097W-A terminator with TDH3 promoter), compared with up to 32-fold decrease (Fig 5E: MMF1 terminator with AMF1 promoter). Similarly as with promoters, strong and weak terminators (100 top and bottom sorted constructs) displayed stronger changes in the opposite directions of expression, as weak ones were expressed overall 2.3-fold more highly (Rank sum test *p*-value < 1e-16), and strong ones overall 2.4-fold more lowly (Rank sum test *p*-value < 1e-16) than the native sequence (Fig S5-1), thus presenting many possibilities for gene expression engineering (Fig 5F). Published experiments confirmed the computationally predicted variability with the ADH1 terminator ^39^, as we measured a strong correlation (Pearson’s *r* = 0.625, *p*-value < 1e-16) to the variability of the fluorescence intensities with the 625 tested promoters (Fig 5G).

We considered that changing the promoters and terminators could have affected gene expression predictions based either on (i) the specific regulatory signals present in the DNA (Fig 3F and 4H) or (ii) the general sequence properties, such as GC content and di-nucleotide composition ^62,63^. We therefore evaluated the effect of removing high-order sequence information (ie. regulatory grammar) by randomly shuffling the regulatory DNA whilst preserving dinucleotide frequencies ^64^. On average, the native DNA regions demonstrated a significant (Levene’s test *p*-value < 1e-16) 88% larger effect on expression variation and a 13.5-fold higher dynamic range compared to randomly shuffled sequences with the same nucleotide composition (Fig S5-2). Accordingly, random sequences could only increase the expression level up to 4.9-fold (Fig S5-2: YIL102C-A promoter with a shuffled variant of its terminator) and decrease it up to 3.2-fold (Fig S5-2: NCE102 promoter with a shuffled variant of its terminator) compared to the native expression level. The observed increase of expression signal by combining native regions with their adjacent counterparts indicated that the correct progression of gene expression requires the presence of a specific regulatory grammar, similar to the one detected above (see Fig 4H), which can be lost when artificially manipulating or carelessly combining the sequences.

## 3. Discussion

In the present study we asked the question: to what extent are gene expression levels encoded in the coding and *cis*-regulatory DNA regions of the gene regulatory structure? This question followed our observation that with all known biological variation of gene expression, based on a collection of RNA-Seq experiments across virtually all to-date tested conditions, 79% of genes were merely within a ± 1-fold change of their median expression levels in ⅔ of the conducted experiments (Fig 1B). By training deep neural networks on the sequences from the entire gene regulatory structure (Fig 1D), we demonstrated that most gene expression levels in *Saccharomyces cerevisiae* are predictable using only DNA encoded information (Fig 1F: *R*^*2*^*_test_* = 0.822). Therefore, 4 orders of magnitude of the transcriptional repertoire can be directly determined from the DNA sequence, irrespective of the experimental condition. Of course, this statement does not object the reality that regulation of gene expression can by highly dynamic between different conditions ^9,39^. However, for the majority of genes, the biological variation is negligible in comparison to the magnitude of expression levels between the genes (Fig 1A), meaning that genes which are highly expressed in the majority of conditions will likely always (79% of protein coding genes) be highly expressed and vice-versa. These differences in gene expression levels are encoded in the DNA and can be read using machine learning. Accordingly, similar results were obtained in 6 other model organisms spanning all kingdoms of life (including bacteria and higher eukaryotes, such as fruit fly, zebrafish, mouse, and human), that covered the complete range of genetic regulatory complexity ^41^, as measured by coding gene density (Table S1-1).

Since individual coding and non-coding parts of the gene regulatory structure (Fig 1D) all play a crucial role in the regulation of gene expression ^9^, we inquired which particular regions contained the highest amounts of information on gene expression levels. While codon frequencies were highly informative about mRNA levels (Fig 2B), we observed that a similar amount of information was encoded in the flanking regions (Fig 2A). Indeed, this was further supported by the result that deep neural networks could predict the codon usage of a gene merely based on that gene’s regulatory sequence (Fig 2C,D). Although the effect of codon usage on overall transcription has been widely studied and debated ^65–67^, their surprisingly strong effects on transcription, also supported by our results (Fig 2B), have only recently started coming to light. The hypothesized mechanisms through which codons affect transcription are: (i) effects on nucleosome positioning ^37^, (ii) premature termination of transcription, usually by mimicking poly-A signals ^68^, or (iii) mRNA toxicity ^69^. Generally, the differences in codon usage between bacterial species can be explained from the dinucleotide content in their non-coding regions ^70^ and it is widely assumed that most inter-species variation in codon usage is attributed to mutational mechanisms ^65,71^. Within a genome, however, since codon usage can be predicted from non-coding regions (Fig 2D), and both coding and regulatory regions are similarly predictive of gene expression levels (Fig 2B), it is likely that the entire gene regulatory structure undergoes a common selection pressure. Accordingly, mutation rates in orthologs from 14 yeast species supported this notion of coevolution between coding and non-coding regions (Fig 2E,F) and showed that the whole gene regulatory structure is a single co-evolved unit. However, despite the ensuing information overlap among the different regions (Fig 2B,D: up to 58%), each region additively contributed to mRNA level prediction (Fig 2A,B) and the entire gene regulatory structure was important for defining over 82% of the gene expression variability (Fig 1F).

The multiple regulatory elements of the gene regulatory structure each control, individually or jointly, the different DNA processing phases required for mRNA transcription, including nucleosome positioning, mRNA synthesis, and mRNA maturation and decay ^9^ (Fig S1-1A). To control enzyme interactions, the DNA therefore contains a plethora of statistically identifiable DNA motifs ^53,54^. The question though remains, which of those motifs are relevant signals for regulating mRNA levels? For instance, based on currently identified JASPAR motifs ^53^, a significant enrichment of known promoter TFBS was found particularly in highly expressed genes (Fig S3-8). However, such an analysis does not uncover which motifs or combinations of motifs a predictive model of mRNA abundance (Fig 1E) finds important. To resolve this we opened the neural network “black box” (Fig 1E, Fig S3-1) and determined the DNA sequences that were causing significant neural network responses (Fig 3A). Although thousands of DNA motifs were uncovered across all the regulatory regions (Fig 3D,F), individual motifs could not explain the entire dynamic range of gene expression, i.e. the same motifs were found in lowly as well as in highly expressed genes (Fig 4A, Fig S4-1). However, by analysing the statistical interactions between motifs, we could identify a much greater indication of expression levels than with single motifs. 9,962 combinations of 2 to 6 motifs were found to co-occur more frequently together than alone across the gene regions (Fig 4C) and were informative of almost the entire dynamic range of expression (Fig 4A, B, F). Moreover, the motif co-occurrence rules also defined more specific ranges of codon usage than single motifs (Fig S4-5), further supporting our results that the entire gene regulatory structure (Fig 1D), including both coding and non-coding regions, is a single co-evolved interacting unit (Fig 2D, E, F).

Finally, we demonstrated that deep neural networks can learn the complex regulatory ‘grammar’ of gene expression directly from an organism’s genome (Fig 1F,H), without any prior knowledge of genetic regulation or the need to perform laborious high-throughput screening experiments with synthetic constructs. Despite the fact that the machine learning model had never seen the synthetic DNA data, it was able to successfully recapitulate fluorescence readouts from published experimental studies ^39,40^ and demonstrate a strong agreement between model predictions and experimental measurements (Fig 1G, Fig S1-5). Furthermore, since the trained models encapsulate the whole interacting regulatory DNA grammar that must be present to correctly drive expression, they can be used to enhance experimental techniques and improve control over gene expression in synthetic biology. We show that the standard approach of introducing a variety of terminators (or promoters) in combination with strong promoters (or terminators) can in fact lead to large variability in each direction of the actual measured levels of expression (Fig 5C,F) ^40,60^. Therefore, using the deep learning models, researchers can now test the effects of different combinations of regions to select the ones that achieve a desired level of gene expression (Fig 5B,E). This can potentially greatly decrease experimental noise, accelerate experimental throughput and thus decrease the overall costs of microorganism development in biotechnology ^3^.

## 4. Methods

### 4.1 Data

Genomic data including transcript and open reading frame (ORF) boundaries were obtained from Ensembl (https://www.ensembl.org/index.html) ^72^, with the exceptions of organisms *Saccharomyces cerevisiae* C288, for which data were obtained from the Saccharomyces Genome Database (https://www.yeastgenome.org/) ^59,73^ and additional published transcript and ORF boundaries were used ^74,75^, and *Escherichia coli* K-12 MG1655, where all data were obtained from the RegulonDB database (http://regulondb.ccg.unam.mx/) ^76^ (Table S1-1). Processed raw RNA sequencing Star counts were obtained from the Digital Expression Explorer V2 database (http://dee2.io/index.html) ^31^ and filtered for experiments that passed quality control. Raw mRNA data were transformed to transcripts per million (TPM) counts ^77^ and genes with zero mRNA output (TPM < 5) were removed (Table S1-2). Extraction of coding and regulatory regions based on transcript and ORF boundaries and processing of mRNA counts yielded gene datasets with paired gene regulatory structure explanatory variables (Fig1D, Fig S1-1C) and mRNA abundance response variables (Fig 1A). Prior to modeling (Fig S1-7), the mRNA counts were Box-Cox transformed ^78^ (see lambda parameters in Table S1-4). DNA sequences were one-hot encoded, UTR sequences were zero-padded up to the specified lengths (Fig 1D), codon frequencies were normalized to probabilities, and 8 mRNA stability variables were computed that included: lengths of 5’-UTR, ORF and 3’-UTR regions, GC content of 5’- and 3’-UTR regions, and GC content at each codon position in the ORF. No significant pairwise correlations were found between the variables (Fig S1-4, Table S1-5), except between mRNA counts and ORF lengths, due to the technical normalization bias from fragment-based transcript abundance estimation ^79^. To obtain mRNA counts that were uncorrelated to gene length, the residual of a linear model, based on ORF lengths as the response variable and mRNA counts as the explanatory variable, was used as the corrected response variable (Fig S1-4). By testing whether the introduced correction could potentially remove biological signal associated with gene length, using data from whole molecule RNA sequencing that does not rely on short-read assembly ^80^, no correlation between gene length and its expression was found (Pearson’s *r* = -0.08, *p*-value < 1e-6; Fig S1-8).

### 4.2 Modeling and statistical analysis

#### Supervised shallow methods

The following regression algorithms were used: linear regression, ridge regression, lasso, elastic net, random forest, support vector machines with nested cross-validation, and k-nearest neighbour regression ^81^. To include information from the regulatory DNA sequences in the shallow models, k-mers of lengths 4 to 6 bp were extracted from the regulatory DNA sequences ^82^ and used as additional explanatory variables. Nested cross validations were performed with these algorithms and GridSearchCV using the Scikit–learn package v0.20.3 with default settings. The coefficient of determination was defined as *R*^2^ = 1 - *SS*_*Residual*_ /*SS*_*Total*_ [Eq. 1], where *SS Residual* is the sum of residual squares of predictions and *SS*_*Total*_ is the total sum of squares.

#### Supervised deep methods

Different neural network architectures were tested that combined: (i) 1 to 4 convolutional neural network (CNN) layers ^83^ (see tested parameters in Table S1-6), which included inception layers ^84^ (ii) 1 to 2 bidirectional recurrent neural network (RNN) layers ^85^, and (iii) 1 to 2 fully connected (FC) layers, in a global architecture layout CNN-RNN-FC ^30,86–88^. Training the networks both (i) concurrently or (ii) consecutively, by weight transfer on different variables (regulatory sequences to CNN and RNN, numeric variables to FC), showed that the architecture yielding best results was a concurrently trained CNN (3 layers)-FC (2 layers) ^12,89–91^, which was used for all models. Batch normalization ^95^ and weight dropout ^96^ were applied after all layers and max-pooling ^97^ after CNN layers (Table S1-6). The Adam optimizer ^92^ (Table S1-6) with mean squared error loss function and ReLU activation function ^93^ with uniform ^94^ weight initialization were used. In total, 26 hyper-parameters were optimized using a tree-structured Parzen estimators approach ^98^ at default settings for 1500 iterations ^99,100^. Tests with different sizes of input regulatory sequences showed that whole regions (Fig 1D) resulted in the most accurate models. To assess model predictions by varying either promoter ^39^ or terminator ^40^ regions, input explanatory variables were constructed based on the specified coding (fluorescence reporters codon frequencies) and regulatory regions (variable region combined with specified adjacent regions). The Keras v2.2 and Tensorflow v1.10 software packages were used.

#### Statistical hypothesis testing

For enrichment analysis, gene ontology slim terms ^101,102^ were obtained from the Saccharomyces Genome Database ^59,73^ and published promoter classifications were used ^26,51^. For statistical hypothesis testing, Scipy v1.1.0 was used with default settings.

### 4.3 DNA sequence analysis

#### Analysis of evolutionary rates

Multi-sequence alignments of 3800 genes (each gene divided into separate promotor, 5’- and 3’-UTR, terminator and coding regions) from fourteen fungal species were generated using Mafft v7.407 ^103^ with the Linsi algorithm and default settings. The resulting 19,000 alignments were analysed with Zorro ^104^ to identify regions of high sequence variability and possible misalignment that could have a negative effect on the phylogenetic signal in the overall sequence. After excluding sites with a confidence score ≤ 0.2, each individual alignment was analysed for 1,000,000 generations in MrBayes v3.2.6 ^105^, with the number of substitution types set to one, and a gamma distribution of substitution rates, to obtain the estimated mean substitution rate (alpha) for each dataset.

#### Relevance analysis

To calculate the relevance of different DNA sequences for model predictions, defined as *Relevance* = (*Y* - *Y* _*occluded*_) / *Y* [Eq. 2], where *Y* is the model prediction, an input dataset with sliding window occlusions was used with the deep models to obtain predictions ^49,106^ (Fig S3-1). The window size of the occlusions was set to either: (i) whole regions, to analyze the *relevance* of region combinations and sensitivity analysis or (ii) 10 bp for motif analysis, determined based on the analysis of *relevance* profiles at difference occlusion sizes using the FastDTW alignment method ^107^ and analysis of the distribution of DNA sequence motif sizes in the JASPAR database ^53^ (Fig S3-2). For clustering of *relevance* profiles, the consensus clustering approach ^108^ was used, as implemented in the package ConsensusClusteringPlus v1.48.0 with the method Partition around medoids (pam) set to 50-fold subsampling of 80% of data points and using the Pearson correlation distance. The number of clusters (*k*) was determined at *k* = 4, for which the relative consensus did not increase more than 10% (Fig S3-9).

#### Motif analysis

Extraction of relevant sequences, i.e. those that significantly (standard deviation ≥ 2) affected gene expression prediction, yielded 169,763 sequences that spanned all the analysed genes (*RSD* < 1) (Fig S3-5). To identify regulatory motifs, clustering of the relevant sequences was performed using CD-HIT v4.8.1 ^109,110^, with recommended settings (a *k*-mer size of 4, 5, and 6 was used in correspondence with sequence identity cutoff 0.8, 0.85, and 0.9, respectively) and a cluster size of 5 sequences, based on the amount of recovered sequences and unique motifs. Multiple sequence alignment on the clustered sequences was performed using Mafft v7.407 ^103^, with the Linsi algorithm and default settings. Position weight motifs (PWMs) were processed using the Biopython package v1.73 ^111^ and motif edges were trimmed below a cutoff of 0.2 bits ^112^. Pairwise comparisons of the PWMs across the regulatory regions and comparisons to JASPAR ^53^ (core fungi, non-redundant) and Yeastract ^54^ databases was performed using Tomtom and Meme suite v4.12 ^113,114^, with recommended settings. Motif co-occurrence was analysed using the FP-growth algorithm as implemented in Apache Spark v2.4 ^115^, with default settings accessed through the Python interface, where *support, confidence* and *lift* were calculated as defined in ^58^. The statistical significance level of co-occurring motifs was determined using the chi-squared test ^116^. Python v3.6 and R v3.6 were used throughout.

## Acknowledgements

We thank Victor Garcia for insightful discussions on codon usage, Clara Correia-Melo and Simran Aulakh for commenting on the manuscript. JZ, FB and AZ are supported by SciLifeLab funding. We gratefully acknowledge the NVIDIA Corporation for supporting this research.

