## Supplementary Information for "Gene expression is encoded in all parts of a co-evolving interacting gene regulatory structure"

### Table of contents

Supplementary Figures … p.2 - p.28

Supplementary Tables … p.29 - p.39

Supplementary References … p.40 - p.41

### Supplementary figures

**A.**

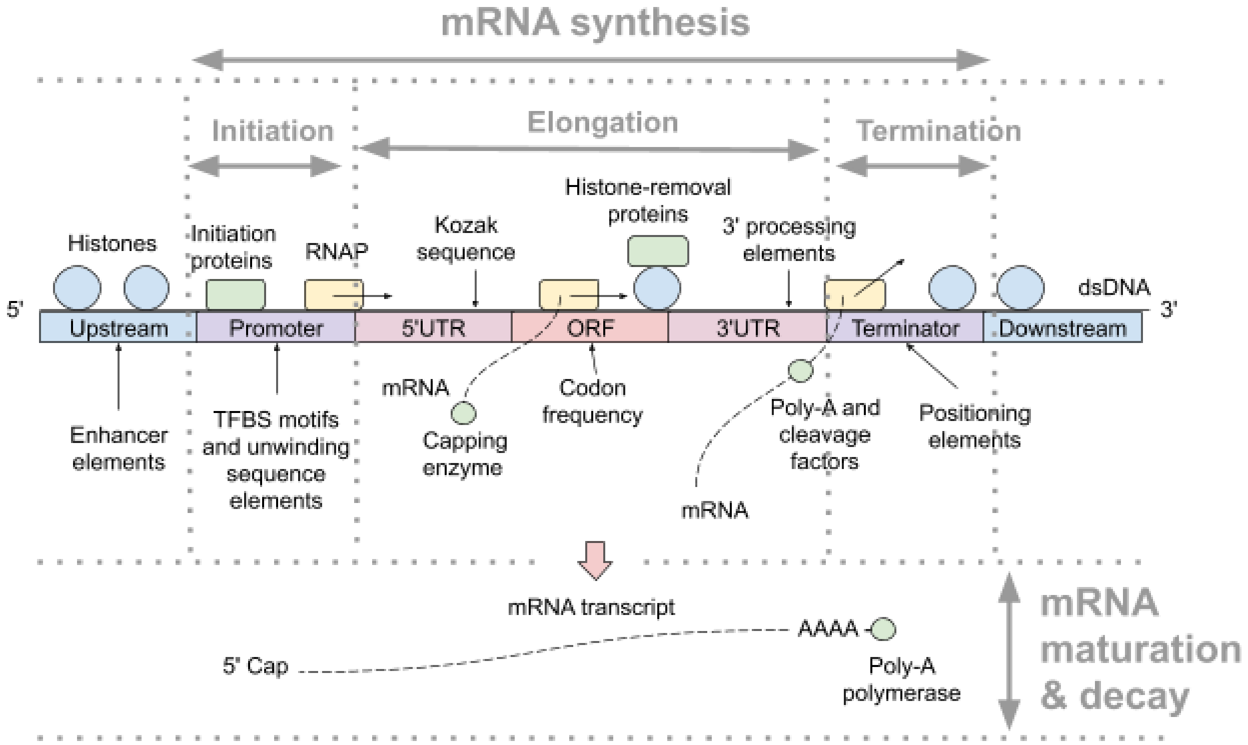


**B.**

| Region | Promoter | 5'UTR | Gene (CDS) | 3'UTR | Terminator |
|---|---|---|---|---|---|
| $\boldsymbol{R^2}$ | 0.46[a] | 0.52[b] | 0.55[c] | >0.16[d] | |

[a] Not genome-wide [1]

[b] Expanded to 0.62 with deep learning [2,3]

[c] Target variable was mRNA half-life, up to 0.59 achieved with extra features [4]

[d] Estimated here based on multiple studies [5,6]

**C.**

| **Region** | **Promoter** | **5'UTR** | **CDS** | **3'UTR** | **Terminator** |
|---|---|---|---|---|---|
| **Regulatory signals** | - Core promoter [7]<br>- TFBS [8]<br>- enhancers [9] | Kozak sequence [2,10] | Codon usage [11,12] | 3' processing elements:<br>- A/T-rich sites [13]<br>- Positioning element [14]<br>- TA-rich efficiency el. [5] | |
| | Nucleosome positioning [6,12,15] | | | | |
| **Size** | 1000 bp | 300 bp | ~300-3000 bp | 350 bp | 500 bp |
| **Positioning** | to TSS | to START [2] | whole | to TTS [13] | from TTS |
| **Data types** | sequence | sequence, variables (2) | variables (67) | sequence, variables (2) | sequence |
| **Sequence data** | yes | yes | no | yes | yes |
| **Variable types** | / | length, GC content [4] | codon freq., length, GC of each wobble pos. [1,16] | length, GC content [4] | / |

**Figure S1-1.** Schematic overview of published knowledge on the gene regulatory structure in *Saccharomyces cerevisiae*. (A) The molecular processes: schematic diagram of mRNA transcription in eukaryotes, detailing separate optimized processes, that form a fine-tuned regulatory system which spans mRNA synthesis, maturation and decay [12]. (B) The information content: overview of the approximate amount of information on gene expression levels that is encoded in each separate region according to published studies. (C) The regulatory system: overview of the known regulatory signals that contain information on gene expression, as well as the sequence parameters and variables used to model and predict gene expression levels in the present study. UTR denotes untranslated regions, ORF open reading frame, CDS coding sequence, TFBS transcription factor binding sites, TSS transcription start site, TTS transcription termination site.

| Pathway | Description | BH adjusted P-value |
|---|---|---|
| GO:0005975 | carbohydrate metabolic process | 5.9e-06 |
| GO:0006091 | generation of precursor metabolites and energy | 2.1e-10 |
| GO:0006520 | cellular amino acid metabolic process | 8.5e-07 |
| GO:0006811 | ion transport | 1e-04 |
| GO:0006865 | amino acid transport | 0.026 |
| GO:0006979 | response to oxidative stress | 0.0021 |
| GO:0008643 | carbohydrate transport | 0.014 |
| GO:0009311 | oligosaccharide metabolic process | 0.002 |
| GO:0032787 | monocarboxylic acid metabolic process | 5.4e-05 |
| GO:0042221 | response to chemical | 0.0082 |
| GO:0045333 | cellular respiration | 8.7e-08 |
| GO:0055085 | transmembrane transport | 0.0065 |
| GO:0055086 | nucleobase-containing small molecule metabolic process | 5.4e-05 |

**Figure S1-2.** Enrichment analysis of gene ontology terms [17,18] in the most variable genes across the entire range of biological conditions (*RSD* > 1).

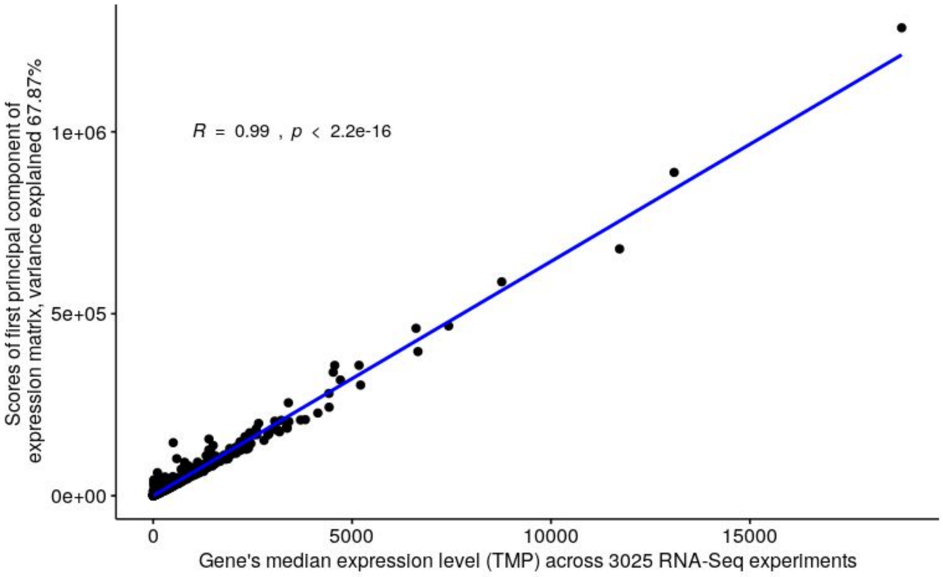


**Figure S1-3.** Median expression levels are representative of a gene's overall expression level across thousands of experiments, based on correlation analysis of the first principal component and median values of the entire matrix of mRNA counts (Pearson's $r = 0.99$, $p$-value < 2e-16). Line denotes least squares fit.

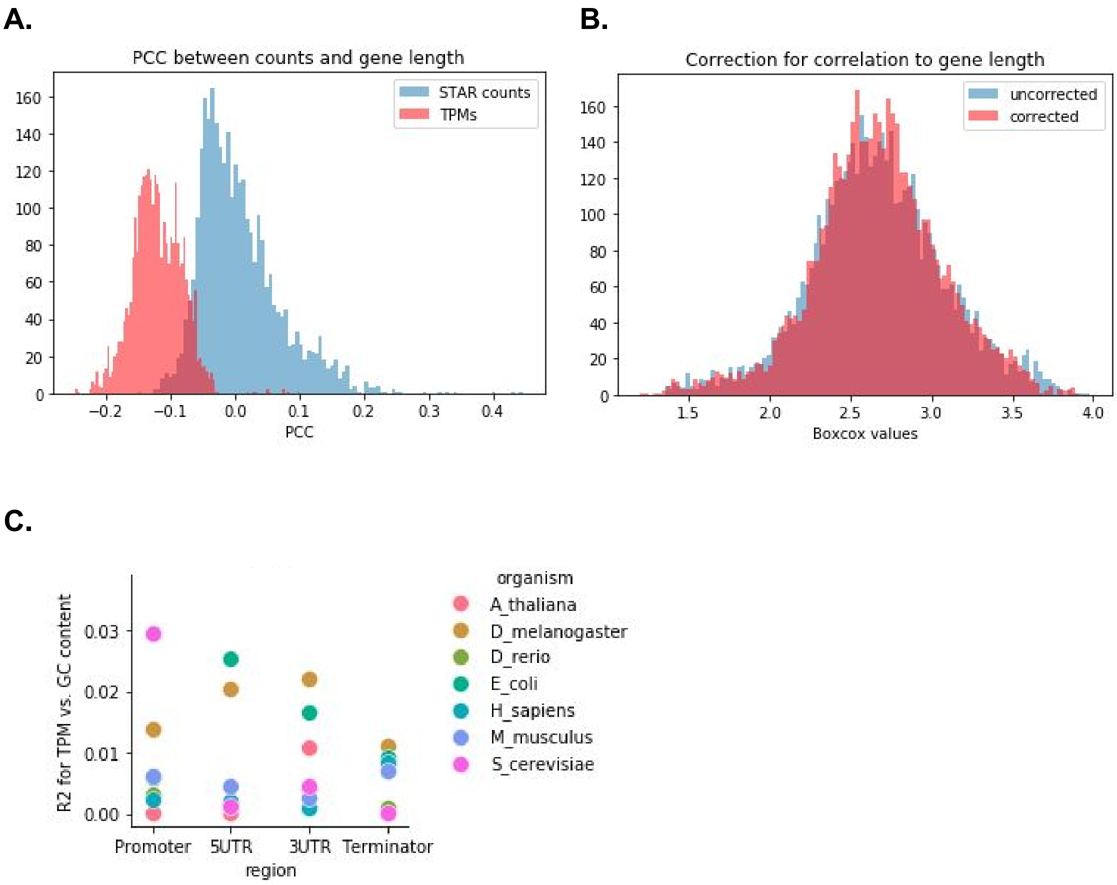


**Figure S1-4.** Overview for RNA-seq data processing with *Saccharomyces cerevisiae*. (A) A detectable level of correlation (above 0.1) was observed between TPM transformed mRNA counts and gene (CDS) length. "PCC" denotes Pearson correlation coefficient. (B) Correction of the TPM target variable, by regressing out gene (CDS) length values, retained all information as the original uncorrected TPM values (Pearson's $r$ = 0.96, $p$-value < 1e-16). (C) Overall GC content of regulatory regions was not predictive of gene expression levels, as the coefficient of determination ($R^2$) between gene expression values and GC content was below 3% for all model organisms.

**A.**

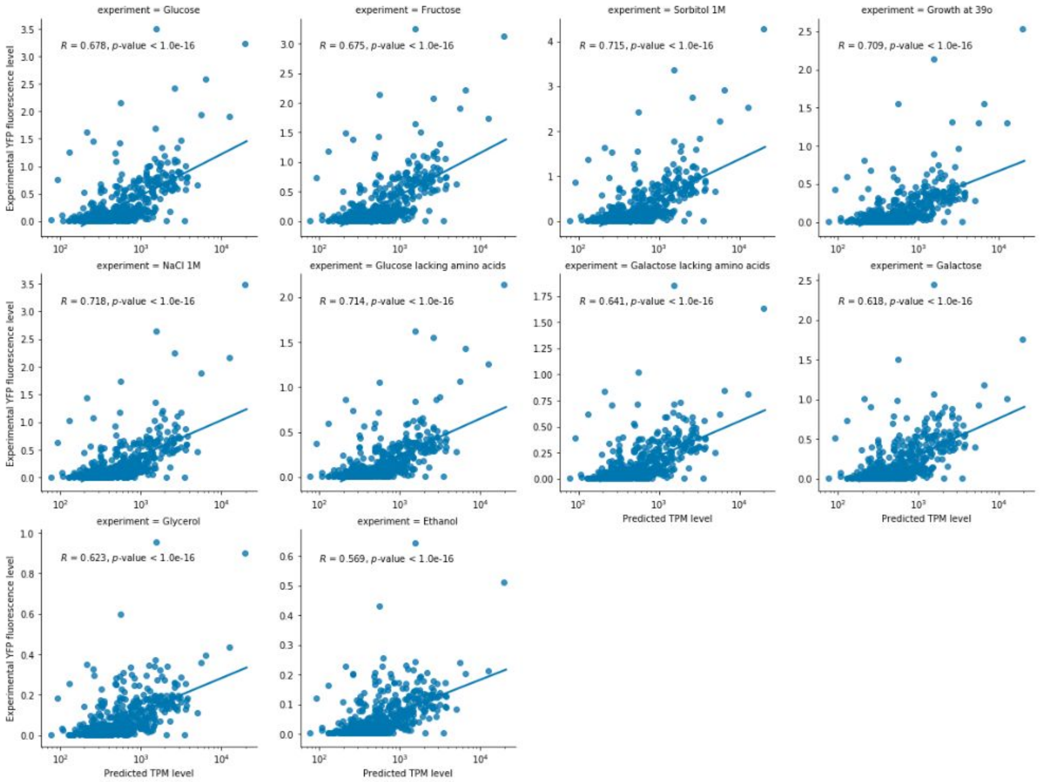


**B.**

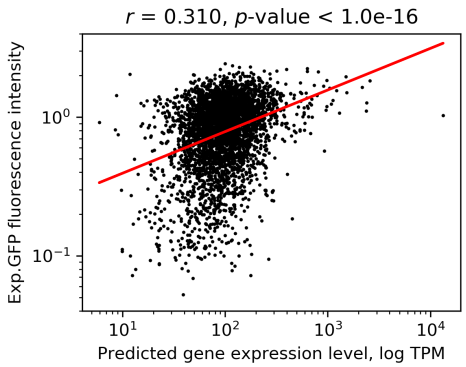


**Figure S1-5.** Model predictions are highly correlated with published experiments. (A) Experimental fluorescence measurements [19] versus predicted expression levels across 10 conditions. (B) Experimental fluorescence measurements [20] versus predicted expression levels. All lines denote least squares fit.

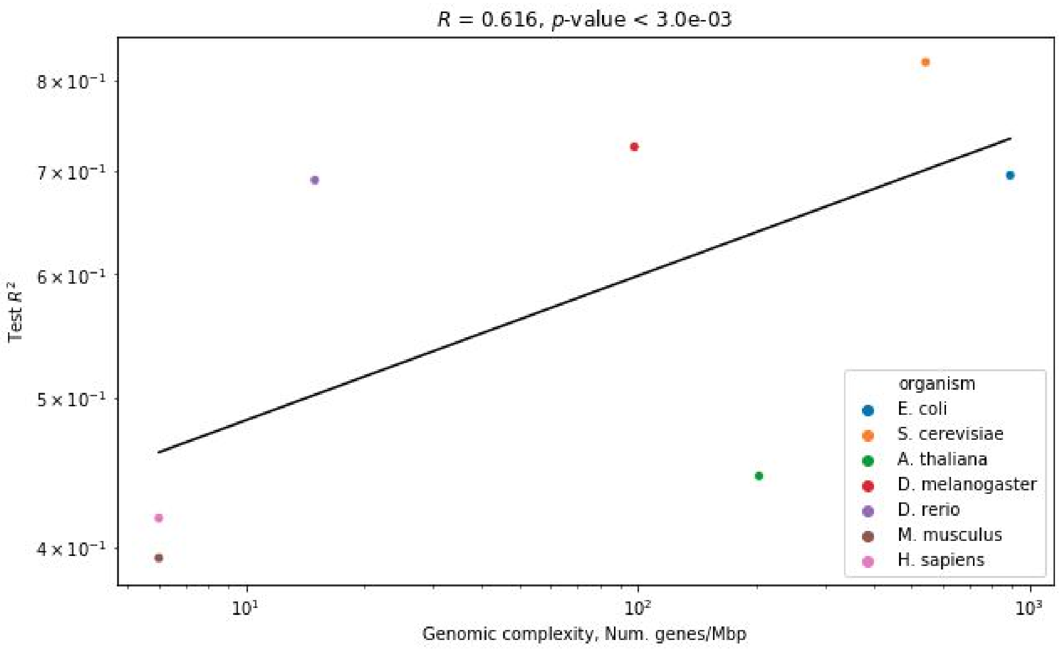


**Figure S1-6.** Correlation analysis between the predictive accuracy of deep learning models ($R^2_{test}$) and the genomic complexity of all model organisms. Line denotes least squares fit.

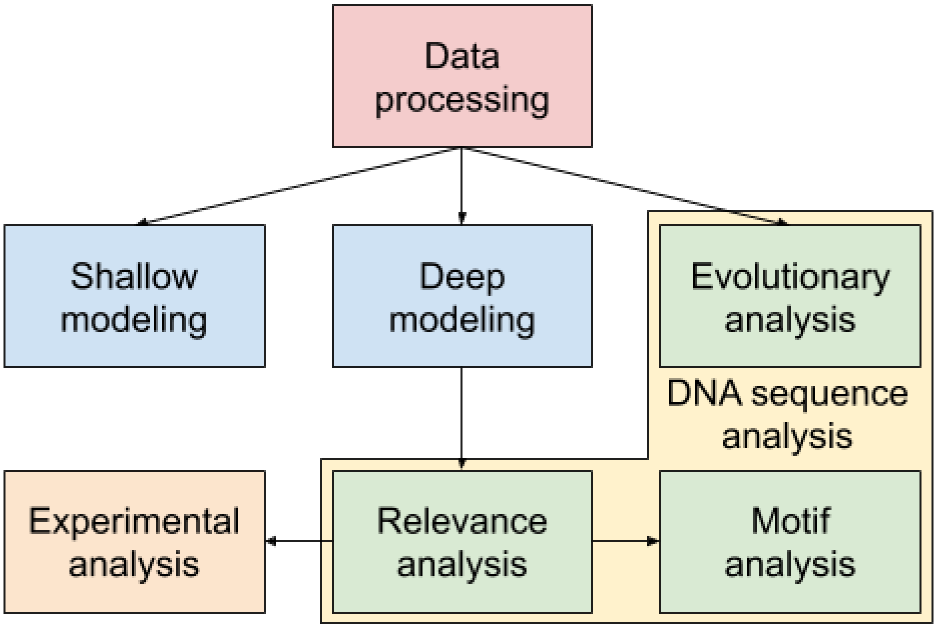


**Figure S1-7.** Overview of computational and experimental pipelines.

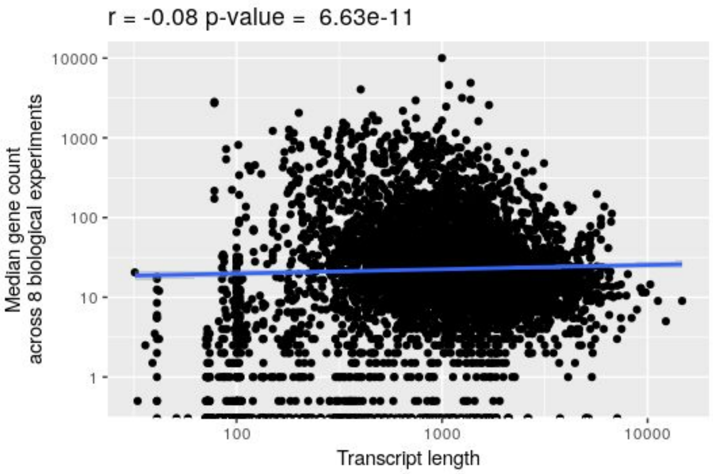


**Figure S1-8.** Correlation analysis between the gene length and the median expression level across experiments per gene, using data from whole molecule RNA-seq with the Oxford Nanopore MinION [21]. Line denotes least squares fit.

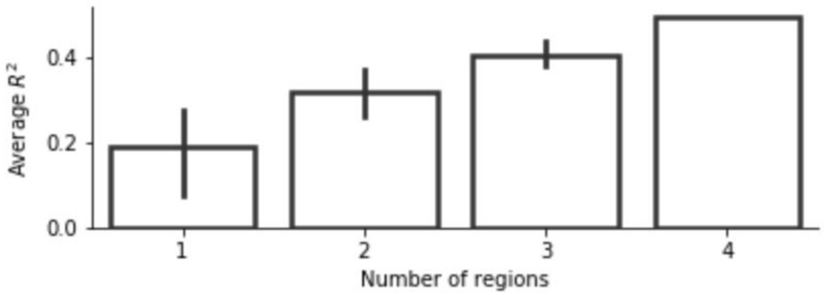


**Figure S2-1.** Effect of combinations of *cis*-regulatory regions on prediction of gene expression levels. The mean and 95% confidence intervals of $R^2_{test}$ at different amounts of regulatory regions are shown.

**A.**

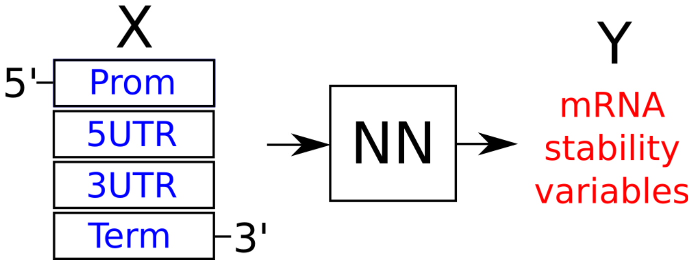


**B.**

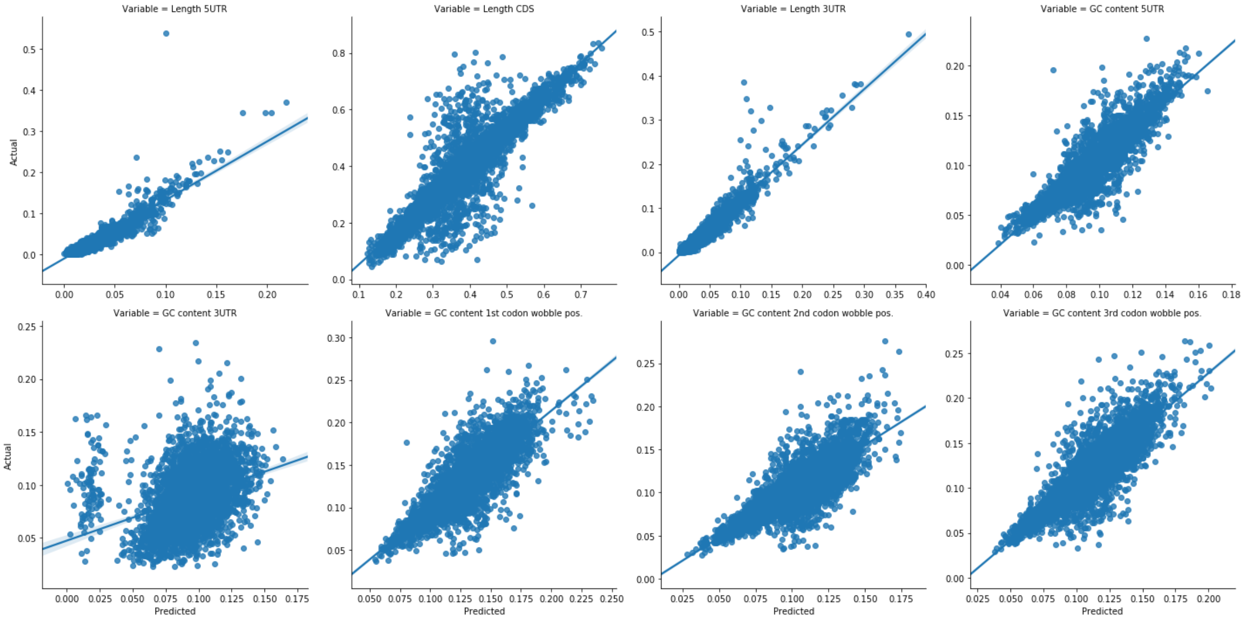


**Figure S2-2.** A CNN was built (A) that could predict nearly 80% of the variation of mRNA stability variables based on input regulatory sequences ($R^2_{test}$ = 0.78). (B) Plots of actual versus predicted stability variables are shown, with individual $R^2_{test}$ values of 0.788, 0.782, 0.864, 0.738, 0.146, 0.682, 0.645 and 0.684, respectively. All lines denote least squares fit.

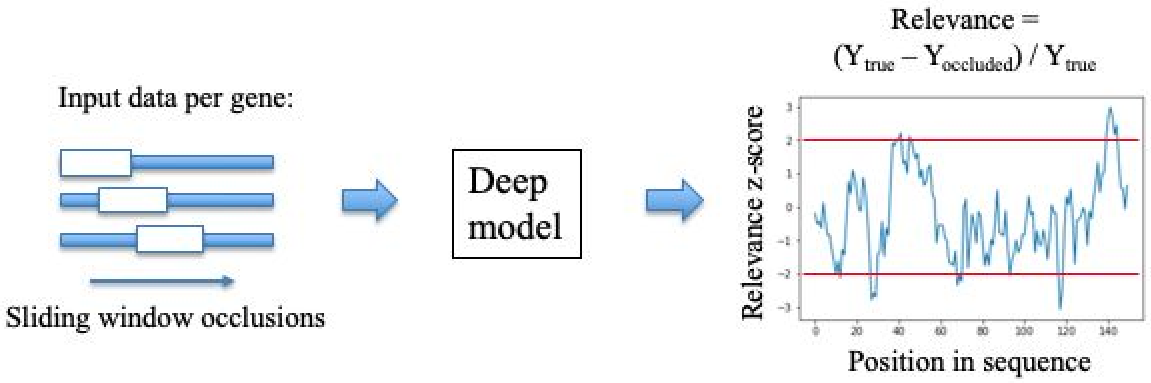


**Figure S3-1.** Schematic overview of the implemented occlusion relevance approach [22,23].

**A.**

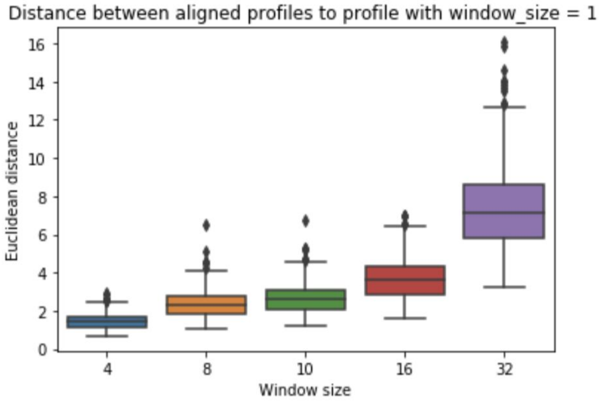


**B.**

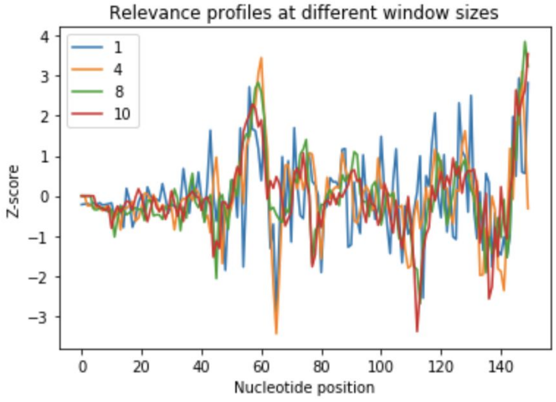


**C.**

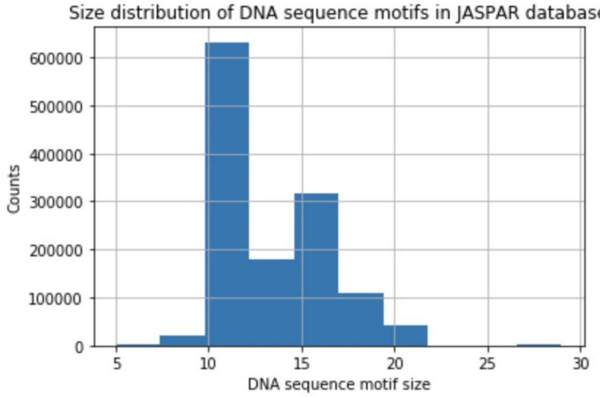


**Figure S3-2.** Analysis of different occlusion window sizes. (A) Euclidean distance between aligned profiles of sizes larger than 1 to the profiles with window size 1. FastDTW alignment method used [24]. (B) An example of the relevance profile with 150bps of a specific promoter region at different window sizes. (C) Size distribution of DNA sequence motifs in JASPAR database (sites file: [http://jaspar.genereg.net/download/sites.tar.gz](http://jaspar.genereg.net/download/sites.tar.gz)). Considering that over 98% of DNA sequence motifs are 10 bps or larger, the analysis suggested that a window size of 10 was a good choice to recover the relevance of true DNA sequence motifs, whilst retaining the relevant information obtainable with the smaller window sizes.

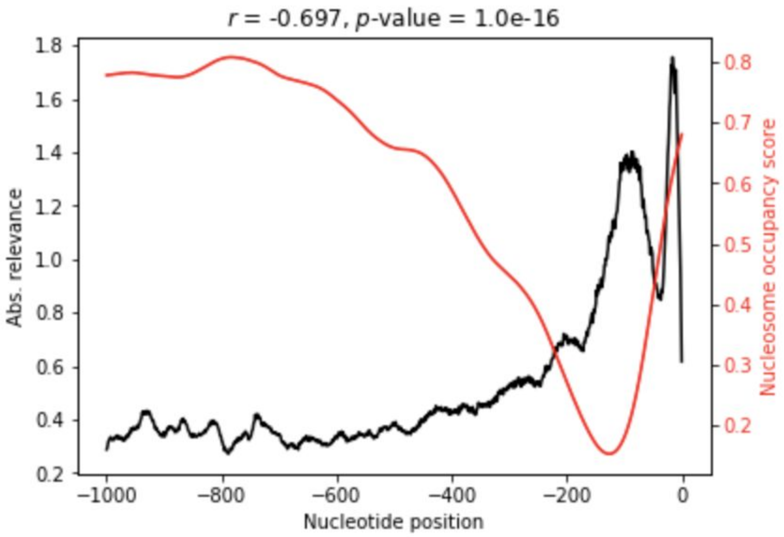


**Figure S3-3.** Strong correlation of absolute relevance in promoter regions and published nucleosome occupancy scores [25] for TFIID regulated genes [26], which were enriched (Fisher's exact test *p*-value < 1e-16) in the *S. cerevisiae* dataset.

| Cluster | Pathway | Description | BH adjusted P-value |
|---|---|---|---|
| 1 | GO:0007059 | chromosome segregation | 1.9e-04 |
| 1 | GO:0033043 | regulation of organelle organization | 4.4e-03 |
| 1 | GO:0048285 | organelle fission | 4.5e-03 |
| 4 | GO:0002181 | cytoplasmic translation | 0.0e+00 |
| 4 | GO:0005975 | carbohydrate metabolic process | 8.1e-04 |
| 4 | GO:0006091 | generation of precursor metabolites and energy | 3.5e-05 |
| 4 | GO:0006414 | translational elongation | 3.1e-06 |
| 4 | GO:0006520 | cellular amino acid metabolic process | 6.8e-05 |
| 4 | GO:0032787 | monocarboxylic acid metabolic process | 7.0e-06 |
| 4 | GO:0051186 | cofactor metabolic process | 6.9e-05 |
| 4 | GO:0055086 | nucleobase-containing small molecule metabolic process | 1.0e-05 |

**Figure S3-4.** Enrichment analysis of gene ontology terms [17,18] in Cluster 4 (with high expressed genes) of the clustered relevance profiles.

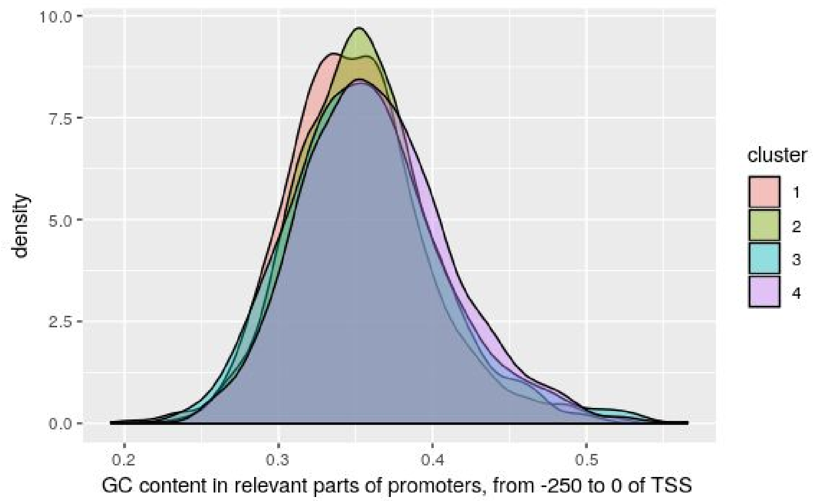


**Figure S3-5.** Clusters of relevance scores are independent of the DNA nucleotide composition.

**A.**

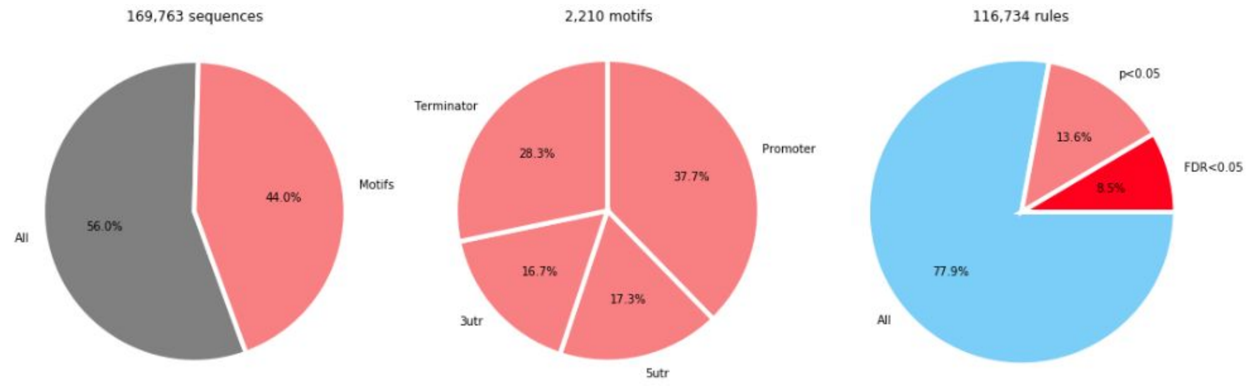


**B.**

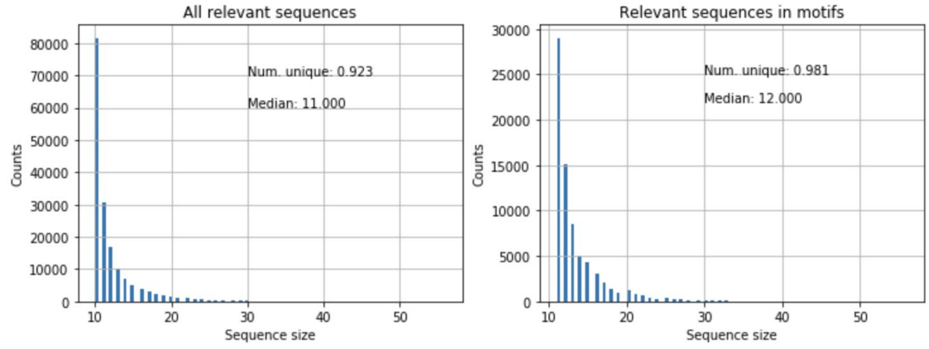


**C.**

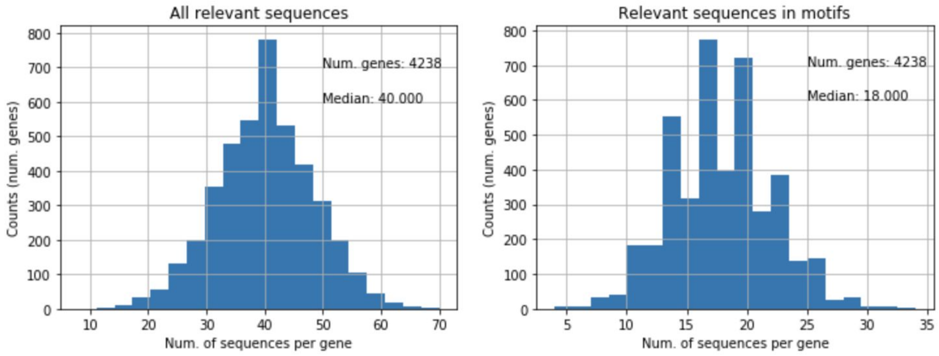


**Figure S3-6.** Analysis of significantly relevant DNA sequences. (A) 169,763 DNA sequences with significant relevance scores (exceeding 95% of range of values, i.e. ± 2 standard deviations) were extracted from the relevance profiles and used to construct regulatory DNA motifs and motif co-occurrence rules. Motif distributions across the *cis*-regulatory regions are shown. (B) Distribution of sizes of all relevant sequences and only those used for constructing the motifs (74,728 at 80% sequence identity cutoff, see Table S3-2). (C) Similarly, distribution of the amount of relevant sequences per gene showed good coverage of the whole set of genes with the extracted regulatory DNA motifs.

**A.**

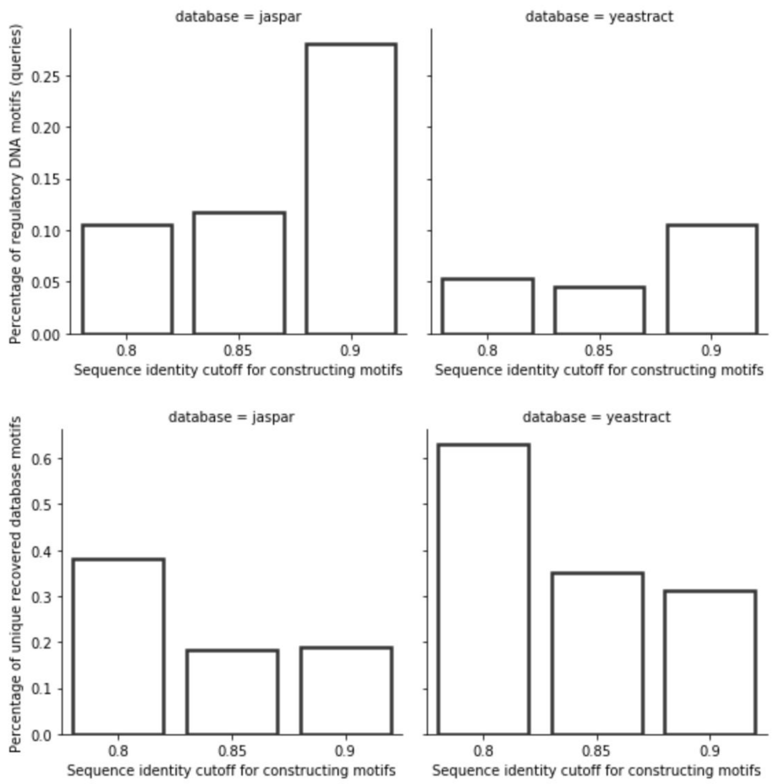


**B.**

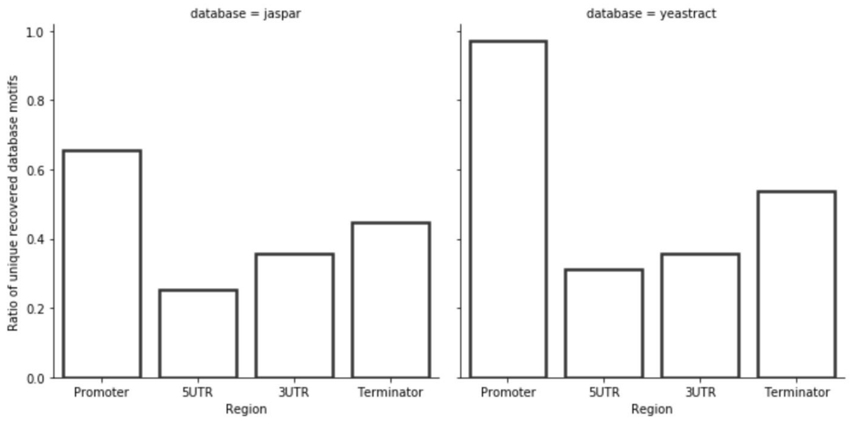


**Figure S3-7.** Comparison of constructed regulatory DNA motifs to JASPAR [27] and Yeastract [28] databases. (A) Although the number of regulatory DNA motifs that are significantly (BH adj. *p*-value < 0.05) similar to ones in databases increases with the increasing sequence identity cutoff used to construct the motifs, the number of unique recovered database motifs decreases, with the exception at the sequence identity cutoff of 0.85 with the Yeastract database. (B) Distribution of significant (BH adj. *p*-value < 0.05) motif hits from the JASPAR [27] and Yeastract [28] databases according to the regulatory regions, where the constructed regulatory motif queries were obtained.

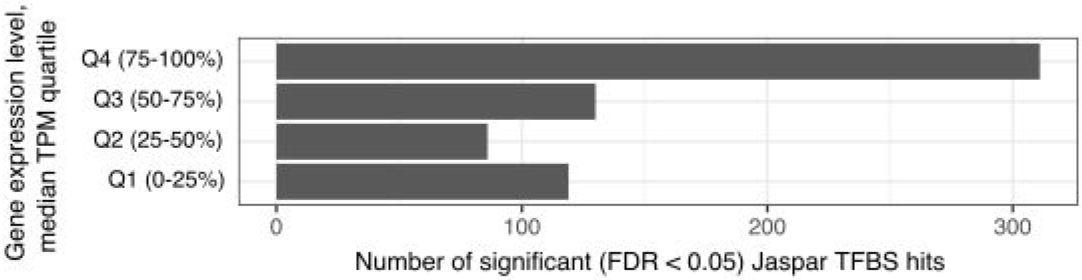


**Figure S3-8.** Enrichment of known yeast TFBS from the Jaspar database [27] in promoters of *Saccharomyces cerevisiae* genes, which were binned into quartiles based on median expression levels.

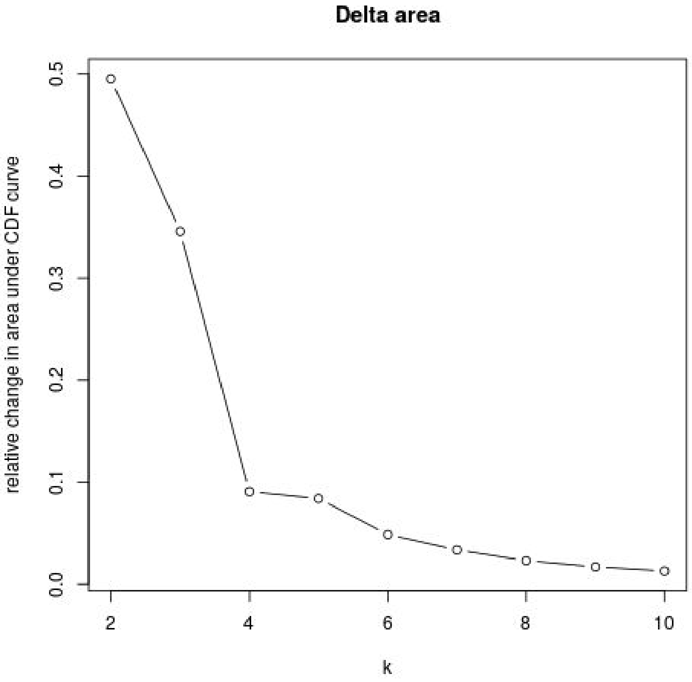


**Figure S3-9.** For clustering of relevance profiles the optimal amount of clusters *k* was determined at 4 (Methods).

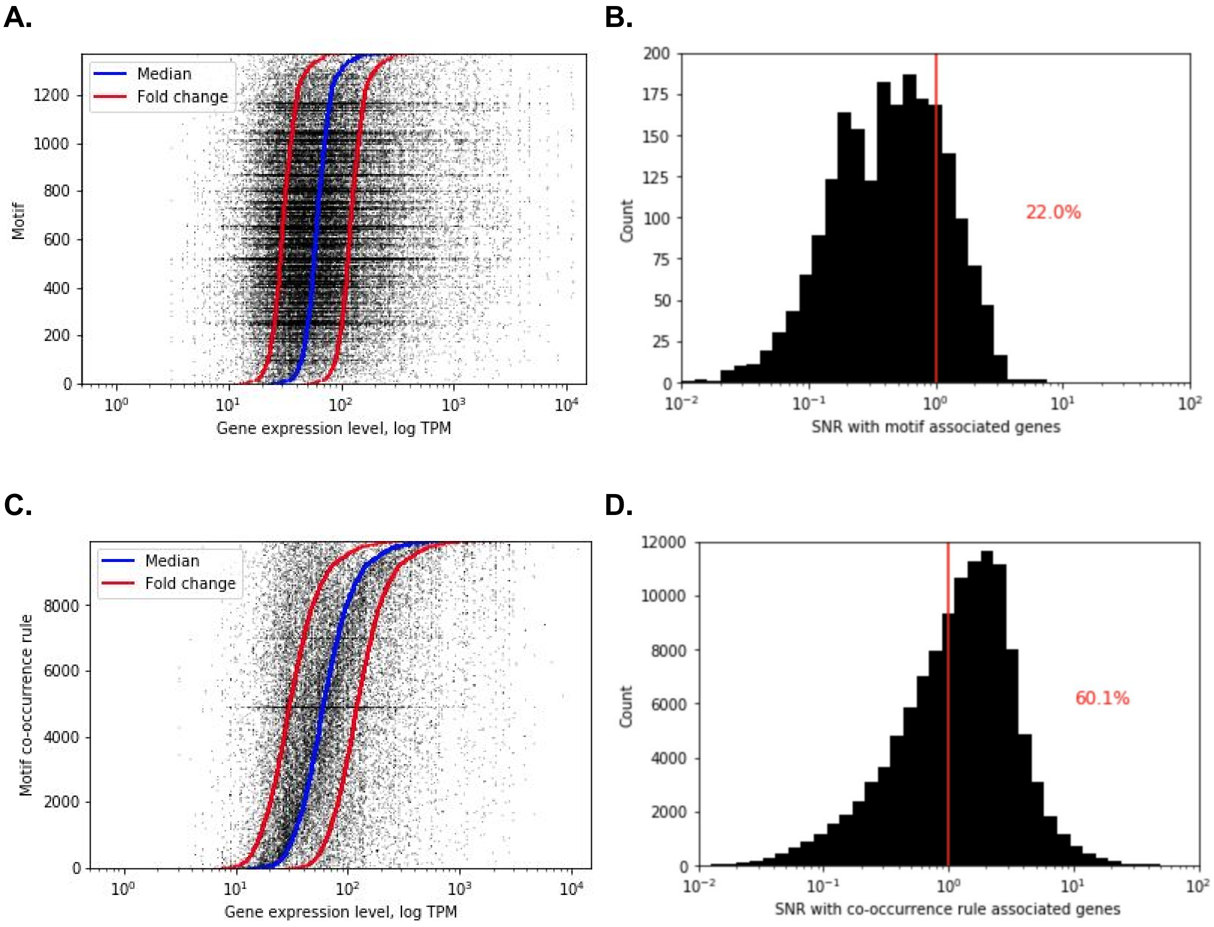


**Figure S4-1.** The range and precision of gene expression regulation with regulatory DNA motifs and motif co-occurrence rules. (A) Expression levels of genes associated with single motifs. (B) Distribution of the signal-to-noise ratio (*SNR*) of expression levels of genes associated with single motifs. Red line denotes an *SNR* of 1. (C) Expression levels of genes associated with motif co-occurrence rules. (D) Distribution of the signal-to-noise ratio (*SNR*) of expression levels of genes associated with motif co-occurrence rules. Red line denotes a *SNR* of 1.

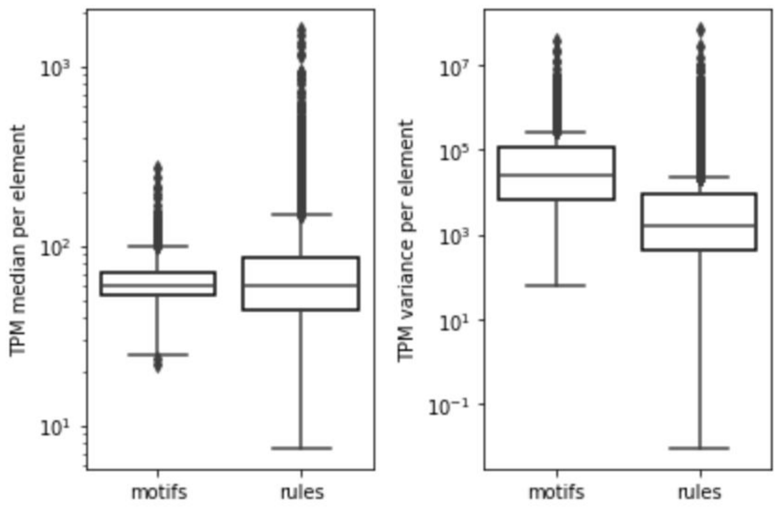


**Figure S4-2.** Median and variance of gene expression levels with genes associated with single motifs or motif co-occurrence rules.

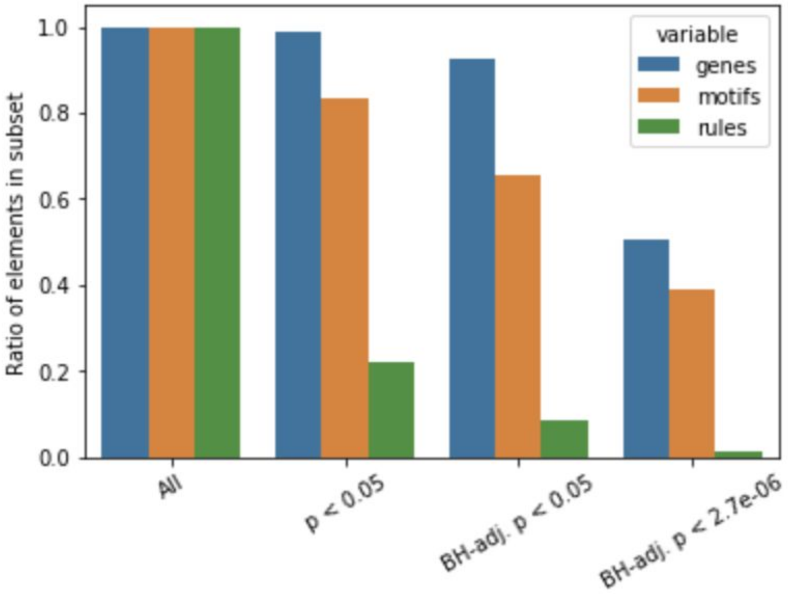


**Figure S4-3.** Ratio of retained elements: unique genes, motifs and rules, with increasing statistical stringency (Fig 4B).

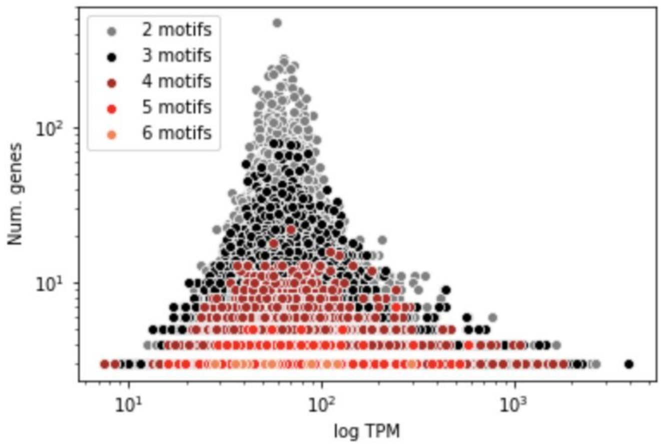


**Figure S4-4.** The number of co-occurring motifs and the amount of genes in a rule versus the average expression level across genes defined by that rule.

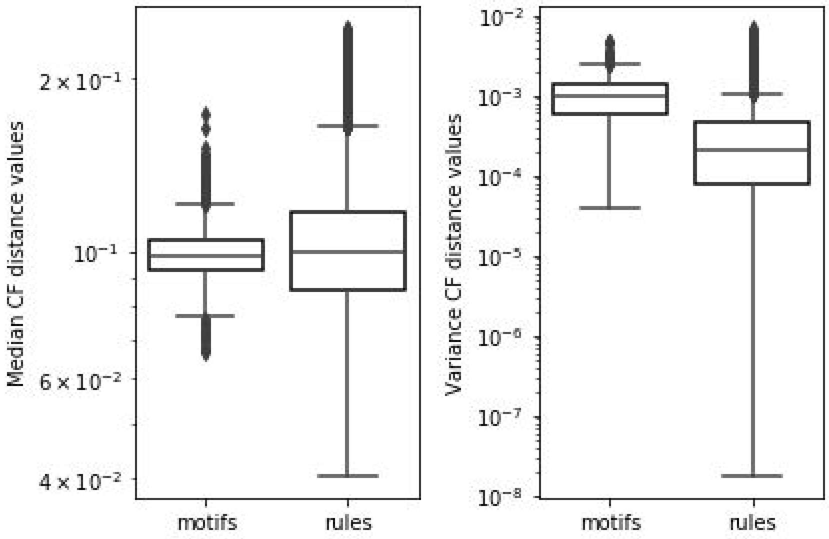


**Fig S4-5.** Median and variance of Euclidean distances between codon frequencies within genes defined by single motifs or motif co-occurrences.

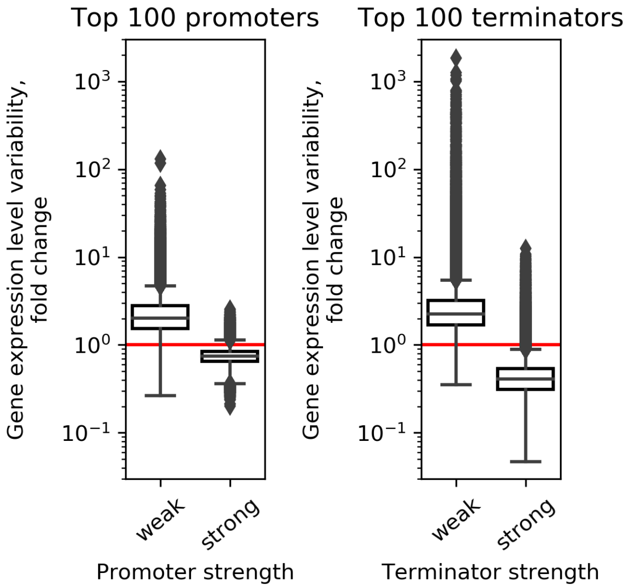


**Figure S5-1.** Variation of gene expression with strong and weak regulatory regions, represented by the selection of 100 top and bottom sorted constructs. (A) Native promoters combined with different terminators. (B) Native terminators combined with different promoters.

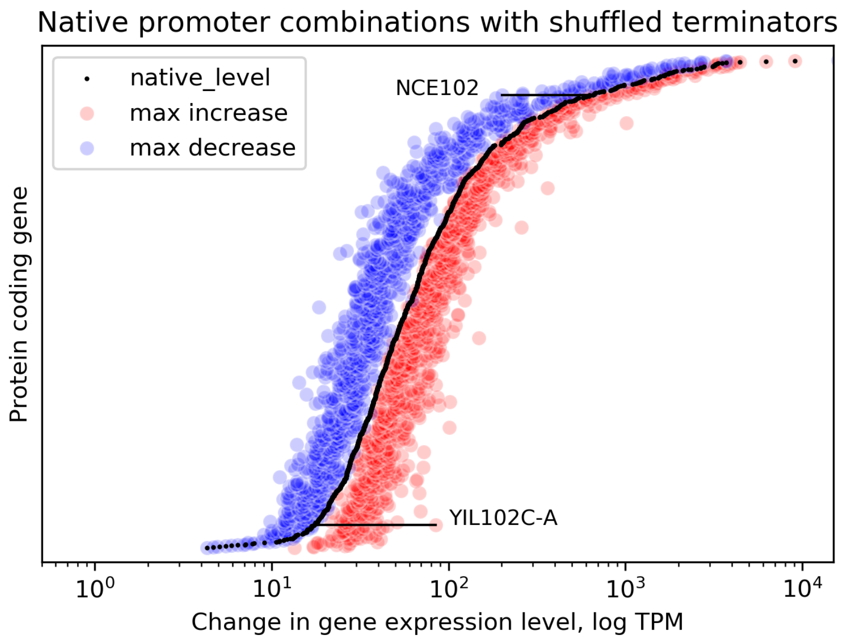


**Figure S5-2.** Evaluation of the effect of removing high-order sequence information (ie. regulatory grammar) by randomly shuffling the regulatory DNA whilst preserving dinucleotide frequencies (Altschul and Erickson 1985). On average, these constructs achieved a 1.4 -fold change in either direction of expression levels and a dynamic range below 1 order of magnitude (6.3 -fold range with YIL102C-A).

### Supplementary Tables

**Table S1-1.** Overview of data and genomic features across the model organisms.

| Organism | Common name | Num. coding genes | Genome size (bps) | Coding gene density | Num. RNAseq datasets used | Num. genes with all regions available |
|---|---|---|---|---|---|---|
| *E. coli* | Bacteria | 4,140 | 4,641,652 | 892 | 355 | 2665 |
| *S. cerevisiae* | Yeast | 6,600 | 12,157,105 | 543 | 3025 | 5112 |
| *A. thaliana* | Plant | 27,655 | 135,670,229 | 204 | 5602 | 22569 |
| *D. melanogaster* | Fruitfly | 13,931 | 142,573,024 | 98 | 4410 | 13317 |
| *D. rerio* | Fish | 25,592 | 1,674,207,132 | 15 | 1084 | 17526 |
| *M. musculus* | Mouse | 22,604 | 3,486,944,526 | 6 | 2365 | 20244 |
| *H. sapiens* | Human | 20,465 | 3,609,003,417 | 6 | 4282 | 18016 |
| Total | / | 120,987 | / | / | 21,123 | 99,449 |
| Average | / | 17,284 | 1,295,028,155 | 252 | 3,018 | 14,207 |

**Table S1-2.** Overview of RNA-seq data across the model organisms.

| **Organism** | **Num. active genes TPM_median > 5** | **Num. genes RSD < 3** | **Num. genes RSD < 2** | **Num. genes RSD < 1** |
|---|---|---|---|---|
| *E. coli (K12)* | 2,154 | 2,012 | 1,737 | 932 |
| *S. cerevisiae* | 4,975 | 4,917 | 4,804 | 4,238 |
| *A. thaliana* | 13,814 | 13,737 | 13,510 | 11,719 |
| *D. rerio* | 7,173 | 7,050 | 6,719 | 4,686 |
| *D. melanogaster* | 9,772 | 9,643 | 9,227 | 5,297 |
| *M. musculus* | 9,951 | 9,785 | 9,370 | 6,585 |
| *H. sapiens* | 9,437 | 9,308 | 8,893 | 6,279 |
| Total | 57,276 | 56,452 | 54,260 | 39,736 |
| Average | 8,182 | 8,065 | 7,751 | 5,677 |
| Relative all | 0.644 | 0.979 | 0.947 | 0.665 |
| Relative Prokarya | 0.808 | 0.934 | 0.806 | 0.433 |
| Relative Yeast | 0.973 | 0.988 | 0.966 | 0.852 |
| Relative Eukarya | 0.616 | 0.987 | 0.951 | 0.704 |

**Table S1-3.** Results of deep modeling across the model organisms.

| **Organism** | ***RSD*** **cutoff** | **Box-Cox lambda** | **Train** $R^2$ | **Validation** $R^2$ | **Test** $R^2$ |
|---|---|---|---|---|---|
| *E. coli* | *2* | -0.147 | 0.778 | 0.645 | 0.695 |
| *S. cerevisiae* | *1* | 0.220 | 0.841 | 0.87 | 0.822 |
| *A. thaliana* | *1* | 0.200 | 0.532 | 0.424 | 0.445 |
| *D. rerio* | *1* | 0.220 | 0.771 | 0.709 | 0.725 |
| *D. melanogaster* | *1* | 0.270 | 0.753 | 0.699 | 0.69 |
| *M. musculus* | *1* | 0.120 | 0.408 | 0.44 | 0.394 |
| *H. sapiens* | *1* | 0.220 | 0.466 | 0.418 | 0.418 |
| Average | / | / | 0.650 | 0.601 | 0.598 |

**Table S1-4.** Overview of the genomic data resources.

| Organism | Strain | Model webpage | Ensembl web | Assembly |
|---|---|---|---|---|
| *E. coli* | K-12 MG1655 | https://ecocyc.org/ | http://bacteria.ensembl.org/Escherichia_coli_str_k_12_substr_mg1655/Info/Index | ASM584v2 |
| *S. cerevisiae* | S288C | https://www.yeastgenome.org/ | http://fungi.ensembl.org/Saccharomyces_cerevisiae/Info/Index | R64-1-1 |
| *A. thaliana* | | https://www.arabidopsis.org/ | http://plants.ensembl.org/Arabidopsis_thaliana/Info/Index | TAIR10 |
| *D. rerio* | | https://zfin.org/ | http://www.ensembl.org/Danio_rerio/Info/Index | GRCz11 |
| *D. melanogaster* | | http://flybase.org/ | http://www.ensembl.org/Drosophila_melanogaster/Info/Index | BDGP6 |
| *M. musculus* | | http://www.informatics.jax.org/ | http://www.ensembl.org/Mus_musculus/Info/Index | GRCm38 |
| *H. sapiens* | | https://www.ncbi.nlm.nih.gov/projects/genome/guide/human/ | http://www.ensembl.org/Homo_sapiens/Info/Index?db=core | GRCh38 |

**Table S1-5.** Correlations between mRNA stability variables.

| Variable 1 | Variable 2 | Pearson's *r* | *p*-value | $R^2$ |
|---|---|---|---|---|
| len3u | gc_3u | 0.239873 | 1.57E-56 | 0.057539 |
| gc_c1 | gc_c3 | 0.180456 | 2.38E-32 | 0.032564 |
| len_5u | gc_c2 | 0.145716 | 1.51E-21 | 0.021233 |
| gc_5u | gc_c3 | 0.142965 | 8.57E-21 | 0.020439 |
| len_5u | len_cd | 0.119115 | 7.26E-15 | 0.014188 |
| gc_3u | gc_c3 | 0.109343 | 9.50E-13 | 0.011956 |
| len_5u | gc_5u | 0.077692 | 4.11E-07 | 0.006036 |
| gc_c2 | gc_c3 | 0.072511 | 2.30E-06 | 0.005258 |
| gc_c1 | gc_c2 | 0.066578 | 1.44E-05 | 0.004433 |
| gc_3u | gc_c1 | 0.058565 | 1.36E-04 | 0.00343 |
| len3u | gc_c2 | 0.041629 | 6.72E-03 | 0.001733 |
| gc_5u | gc_3u | 0.040962 | 7.65E-03 | 0.001678 |
| len_cd | gc_5u | 0.037118 | 1.57E-02 | 0.001378 |
| gc_5u | gc_c1 | 0.026558 | 8.39E-02 | 0.000705 |
| len_5u | len3u | 0.011822 | 4.42E-01 | 0.00014 |
| gc_5u | gc_c2 | -0.008987 | 5.59E-01 | 0.000081 |
| gc_3u | gc_c2 | -0.015623 | 3.09E-01 | 0.000244 |
| len_5u | gc_3u | -0.017595 | 2.52E-01 | 0.00031 |
| len3u | gc_c1 | -0.021041 | 1.71E-01 | 0.000443 |
| len3u | gc_c3 | -0.032953 | 3.19E-02 | 0.001086 |
| len3u | gc_5u | -0.041471 | 6.93E-03 | 0.00172 |
| len_5u | gc_c3 | -0.04148 | 6.92E-03 | 0.001721 |
| len_cd | gc_c2 | -0.051434 | 8.09E-04 | 0.002646 |
| len_cd | gc_3u | -0.051623 | 7.74E-04 | 0.002665 |
| len_5u | gc_c1 | -0.070484 | 4.37E-06 | 0.004968 |
| len_cd | len3u | -0.079376 | 2.29E-07 | 0.006301 |
| len_cd | gc_c1 | -0.163237 | 1.07E-26 | 0.026646 |
| len_cd | gc_c3 | -0.2974 | 2.71E-87 | 0.088447 |

**Table S1-6.** Hyper-parameters used with deep learning algorithms. CNN denotes convolutional neural networks, RNN recurrent neural networks and FC fully connected neural networks.

| Type | Parameter name | Values | Value range |
|---|---|---|---|
| Global | num epochs | 500 | fixed |
| | early stopping min delta | 0.01 | fixed |
| | early stopping patience | 50 | fixed |
| | LRS* epoch drop | 10 | fixed |
| | learning rate | (0.00001,0.1) | log variable |
| | beta_1 | (0.5,0.95) | uniform variable |
| | beta_2 | (0.9,0.95) | uniform variable |
| | epsilon | 1.00E-07 | fixed |
| | mbatch | [64.128,256] | fixed |
| CNN | kernel size | [10, 20, 30, 40] | fixed |
| | filters | [32, 64, 128] | fixed |
| | dilation | [1, 2, 4] | fixed |
| | stride | 1 | fixed |
| | max-pool size | [1, 2, 4] | fixed |
| | max-pool stride | [1, 2] | fixed |
| | dropout | (0, 1) | uniform variable |
| RNN | kernel size | 64 | fixed |
| | dropout | (0, 1) | uniform variable |
| FC | dense size | [32, 64, 128] | fixed |
| | dropout | (0, 1) | uniform variable |

* Learning rate scheduler

**Table S2-1.** Deep modeling results using different combinations of codon probabilities, mRNA stability variables and regulatory sequences.

| Input variable combinations | Target | Layer type | Input type | Train $R^2$ | Validation $R^2$ | Test $R^2$ |
|---|---|---|---|---|---|---|
| Regulatory regions | TPM | CNN | Sequences | 0.845 | 0.575 | 0.492 |
| mRNA stability | TPM | Dense (FC) | 8 variables | 0.386 | 0.471 | 0.378 |
| Coding regions | TPM | Dense (FC) | 64 variables | 0.715 | 0.742 | 0.69 |
| Regulatory + stability | TPM | Dense (FC) | 72 variables | 0.597 | 0.603 | 0.558 |
| Regulatory + coding | TPM | CNN + Dense | Seq. + 64 vars. | 0.824 | 0.862 | 0.816 |
| Codoning + stability | TPM | Dense (FC) | 72 variables | 0.721 | 0.751 | 0.755 |
| All | TPM | CNN + Dense | Seq. + 72 vars. | 0.841 | 0.87 | 0.822 |
| Regulatory regions | Codon prob. | CNN + Dense | Sequences | 0.538 | 0.543 | 0.582 |
| Regulatory regions | Codon prob. | CNN + Dense | Sequences | 0.969 | 0.776 | 0.779 |

**Table S2-2.** Shallow modeling results using linear regression with different combinations of codon probabilities, mRNA stability variables and kmers of size 4 to 6 as features.

| Features | Kmer size | Train $R^2$ | Test $R^2$ | Train *MSE** | Test *MSE* | Fit time | Score time |
|---|---|---|---|---|---|---|---|
| codon_stability | 4 | 0.699 | 0.685 | 0.039 | 0.040 | 0.030 | 0.002 |
| codon | 4 | 0.693 | 0.681 | 0.039 | 0.041 | 0.037 | 0.002 |
| codon_stability_kmers | 4 | 0.728 | 0.674 | 0.035 | 0.042 | 0.456 | 0.005 |
| codon_kmers | 4 | 0.720 | 0.667 | 0.036 | 0.043 | 0.470 | 0.007 |
| stability_kmers | 4 | 0.265 | 0.159 | 0.094 | 0.108 | 0.325 | 0.005 |
| stability | 4 | 0.147 | 0.142 | 0.109 | 0.110 | 0.002 | 0.001 |
| kmers | 4 | 0.153 | 0.031 | 0.109 | 0.124 | 0.409 | 0.005 |
| codon_stability | 5 | 0.699 | 0.685 | 0.039 | 0.040 | 0.077 | 0.002 |
| codon | 5 | 0.693 | 0.681 | 0.039 | 0.041 | 0.018 | 0.002 |
| codon_stability_kmers | 5 | 0.792 | 0.593 | 0.027 | 0.052 | 7.497 | 0.018 |
| codon_kmers | 5 | 0.788 | 0.585 | 0.027 | 0.053 | 6.992 | 0.016 |
| stability | 5 | 0.147 | 0.142 | 0.109 | 0.110 | 0.002 | 0.001 |
| stability_kmers | 5 | 0.423 | -0.085 | 0.074 | 0.139 | 5.278 | 0.015 |
| kmers | 5 | 0.343 | -0.234 | 0.084 | 0.158 | 6.558 | 0.018 |
| codon_stability | 6 | 0.699 | 0.685 | 0.039 | 0.040 | 0.057 | 0.002 |
| codon | 6 | 0.693 | 0.681 | 0.039 | 0.041 | 0.021 | 0.002 |
| stability | 6 | 0.147 | 0.142 | 0.109 | 0.110 | 0.002 | 0.001 |
| codon_stability_kmers | 6 | 1.000 | -8.008 | 0.000 | 1.150 | 234.612 | 0.060 |
| codon_kmers | 6 | 1.000 | -8.313 | 0.000 | 1.188 | 237.973 | 0.065 |
| stability_kmers | 6 | 1.000 | -15.425 | 0.000 | 2.097 | 230.862 | 0.056 |
| kmers | 6 | 1.000 | -17.296 | 0.000 | 2.333 | 235.376 | 0.068 |

* Mean squared error

**Table S2-3.** 14 yeast species used to analyse co-evolution of regulatory and coding regions.

| Clade [29] | Species | Strain | Ensembl availability | Assembly |
|---|---|---|---|---|
| Saccharomyces | Saccharomyces cerevisiae | S288C | https://fungi.ensembl.org/Saccharomyces_cerevisiae/Info/Index | R64-1-1 |
| Saccharomyces | Saccharomyces eubayanus | FM1318 | http://fungi.ensembl.org/Saccharomyces_eubayanus_gca_001298625/Info/Index | SEUB3.0 |
| | Candida glabrata | CSB 138 | https://fungi.ensembl.org/_candida_glabrata_gca_000002545/Info/Index | ASM254v2 |
| Kluyveromyces | Kluyveromyces lactis | NRRL Y-1140 | http://fungi.ensembl.org/Kluyveromyces_lactis_gca_000002515/Info/Index | ASM251v1 |
| Candida | Candida albicans | SC 5314 | http://fungi.ensembl.org/Candida_albicans_sc5314_gca_000784635/Info/Index | Cand_albi_SC5314_V4 |
| Candida | Debaryomyces hansenii | CBS767 | http://fungi.ensembl.org/Debaryomyces_hansenii_cbs767_gca_000006445/Info/Index | ASM644v2 |
| | Yarrowia lipolytica | | http://fungi.ensembl.org/Yarrowia_lipolytica_gca_900087985/Info/Index | YALIA101 |
| Schizosaccharomyces | Schizosaccharomyces pombe | 972h- | http://fungi.ensembl.org/Schizosaccharomyces_pombe/Info/Index | ASM294v2 |
| Schizosaccharomyces | Schizosaccharomyces japonicus | YFS 275 | http://fungi.ensembl.org/Schizosaccharomyces_japonicus/Info/Index | GCA_000149845.2 |
| | Saccharomyces kudriavzevii | IFO 1802 | http://fungi.ensembl.org/Saccharomyces_kudriavzevii_ifo_1802_gca_000167075/Info/Index | Saccharomyces_kudriavzevii_strain_IFO1802_v1.0 |
| | Saccharomyces arboricola | H-6 | http://fungi.ensembl.org/Saccharomyces_arboricola_h_6_gca_000292725/Info/Index | SacArb1.0 |
| | Saccharomyces sp boulardii | biocodex | http://fungi.ensembl.org/Saccharomyces_sp_boulardii__gca_001298375/Info/Index | ASM129837v2 |
| | Kluyveromyces marxianus | DMKU3 1042 | https://fungi.ensembl.org/Kluyveromyces_marxianus_dmku3_1042_gca_001417885/Info/Index | Kmar_1.0 |
| | Kluyveromyces_dobzhanskii | CBS 2104 | http://fungi.ensembl.org/Kluyveromyces_dobzhanskii_cbs_2104_gca_000820885/Info/Index | KLDO_01 |

**Table S3-1.** Construction of regulatory DNA motifs at different sequence identity cutoffs.

| Seq. id. | Num. motifs | % Relevant sequences in motifs | % Jaspar targets | % Motif overlap between gene regions | Num. co-occuring motifs |
|---|---|---|---|---|---|
| 0.8 | 2210 | 0.4401901474 | 0.318182 | 0.15268 | 116,734 |
| 0.85 | 2786 | 0.2716610804 | 0.284091 | 0.269168 | 12,809 |
| 0.9 | 1152 | 0.08214982063 | 0.210227 | 0.140091 | 408 |

**Table S4-1.** Groups of motif co-occurrence rules with a common Jaspar TFBS motif in promoter regions that define expression levels in an over 30 fold range of values.

| Motif name | BH adj. *p*-value | Regions with differing motifs | Num. rules | Num. genes | Fold change |
|---|---|---|---|---|---|
| NHP6B | 0.00448922 | (3UTR, 5UTR, Promoter, Terminator) | 144 | 144 | 648.016 |
| ABF1 | 0.0499867 | (3UTR, 5UTR, Promoter, Terminator) | 32 | 42 | 298.673 |
| STB3 | 0.000661653 | (3UTR, 5UTR, Promoter, Terminator) | 46 | 64 | 166.031 |
| HAP3 | 0.0358934 | (3UTR, 5UTR, Promoter, Terminator) | 83 | 102 | 132.435 |
| AZF1 | 0.0334191 | (3UTR, 5UTR, Promoter, Terminator) | 5 | 12 | 100.093 |
| CBF1 | 0.0036968 | (3UTR, 5UTR, Promoter, Terminator) | 58 | 65 | 73.3709 |
| CUP2 | 0.000975943 | (3UTR, 5UTR, Promoter, Terminator) | 3 | 21 | 55.2978 |
| CUP9 | 0.022899 | (3UTR, 5UTR, Promoter, Terminator) | 54 | 77 | 53.2898 |
| SFP1 | 0.00201301 | (3UTR, 5UTR, Promoter, Terminator) | 7 | 14 | 51.6828 |
| RSC3 | 0.0423265 | (3UTR, 5UTR, Promoter, Terminator) | 10 | 17 | 42.2857 |
| SUM1 | 0.0384774 | (3UTR, 5UTR, Promoter, Terminator) | 10 | 15 | 35.5945 |
| NSI1 | 0.0178329 | (3UTR, 5UTR, Promoter, Terminator) | 18 | 29 | 34.9994 |
